# PEMA: from the raw .fastq files of 16S rRNA and COI marker genes to the (M)OTU-table, a thorough metabarcoding analysis

**DOI:** 10.1101/709113

**Authors:** Haris Zafeiropoulos, Ha Quoc Viet, Katerina Vasileiadou, Antonis Potirakis, Christos Arvanitidis, Pantelis Topalis, Christina Pavloudi, Evangelos Pafilis

## Abstract

**Background:** Environmental DNA (eDNA) and metabarcoding, allow the identification of a mixture of individuals and launch a new era in bio- and eco-assessment. A number of steps are required to obtain taxonomically assigned (Molecular) Operational Taxonomic Unit ((M)OTU) tables from raw data. For most of these, a plethora of tools is available; each tool’s execution parameters need to be tailored to reflect each experiment’s idiosyncrasy. Adding to this complexity, for such analyses, the computation capacity of High Performance Computing (HPC) systems is frequently required.

Software containerization technologies ease the sharing and running of software packages across operating systems; thus, they strongly facilitate pipeline development and usage. Likewise are programming languages specialized for big data pipelines, incorporating features like roll-back checkpoints and on-demand partial pipeline execution.

**Findings:** PEMA is a containerized assembly of key metabarcoding analysis tools with a low effort in setting up, running and customizing to researchers’ needs. Based on third party tools, PEMA performs reads’ pre-processing, clustering to (M)OTUs and taxonomy assignment for 16S rRNA and COI marker gene data. Due to its simplified parameterisation and checkpoint support, PEMA allows users to explore alternative algorithms for specific steps of the pipeline without the need of a complete re-execution. PEMA was evaluated against previously published datasets and achieved comparable quality results.

**Conclusions:** Given its time-efficient performance and its quality results, it is suggested that PEMA can be used for accurate eDNA metabarcoding analysis, thus enhancing the applicability of next-generation biodiversity assessment studies.

## Background

Metabarcoding inaugurates a new era in bio- and eco-monitoring. However, from the output of a sequencer to an amplicon study analysis results, it takes a long way.

Well-established pipelines are available to process metabarcoding data (mothur [1], QIIME 2 [2], LotuS [3]) for the case of 16S rRNA marker gene and bacterial communities. However, there is none that can be used in a straightforward way for metabarcoding analysis of eukaryotic organisms. For this to be functional, adaptation to other marker genes (e.g COI) is required. Furthermore, the pipelines mentioned above, although entrenched, they still suffer from a series of hurdles: technical difficulties in installation and use, strict limitations in setting parameters for the algorithms invoked, incompetence in partial re-execution of an analysis, are among the most prominent.

Moreover, given the computational demands of such analyses, access to High Performance Computing (HPC) systems, might be mandatory, for example, to process studies with large number of samples. This is rather timely given the ongoing investment of national and international efforts (for example [4]) to serve the broad biological community via commonly accessible infrastructures.

PEMA is an open-source pipeline that bundles state-of-the-art bioinformatic tools for all necessary steps of amplicon analysis and aims to address the issues mentioned above. It is designed for paired-end sequencing studies and is implemented in the BigDataScript (BDS) [5] programming language. BDS’s *ad hoc* task parallelism and task synchronization, supports heavyweight computation which PEMA inherits. In addition, BDS supports *checkpoint* files that can be used for partial re-execution and crash recovery of the pipeline. PEMA builds on this feature to serve tool and parameter exploratory customization for optimal metabarcoding analysis fine tuning. Switching effortlessly between clustering algorithms is a pertinent example. Finally, via the Docker [6] and Singularity [7], the latter HPC-centered, software containerization technologies, PEMA is distributed in an easy to download and install fashion on a range of systems from regular computers, to cloud or HPC environments.

Beyond the technical aspects and from the biology perspective, PEMA supports the metabarcoding analysis of both prokaryotic communities (based on the 16S rRNA marker gene) and eukaryotic ones (based on the COI marker gene).

Two clustering algorithms, Swarm v2 [8] and CROP [9], are employed for the clustering of reads in Molecular Operational Taxonomic Units (MOTUs) in the COI marker gene case. VSEARCH [10] is used for the 16S rRNA gene case. Taxonomy assignment is performed in an alignment-based approach, making use of the CREST LCAClassifier [11] and the Silva database [12]. For the COI marker gene, the RDPClassifier [13] and the MIDORI database [14] are used. In the 16S marker gene case, phylogeny-based assignment is also supported, based on RAxML-ng [15], EPA-ng [16] and Silva, as well as ecological and phylogenetic analysis via the “phyloseq” R package [17].

All the pipeline-controlling and third-party-module parameters are defined in a plain *parameter-value pair* text file. Its straightforward format eases the analysis fine tuning complementary to the aforementioned *checkpoint* mechanism. A tutorial about PEMA and installation guidance can be found on PEMA’s GitHub repository (https://github.com/hariszaf/pema).

## Implementation

PEMA’s architecture comprises four main parts taking place in tandem (Figure 1). Detailed description of the tools invoked by PEMA and their licences is included in Additional file 1: Supplementary Methods.

**Figure 1:**
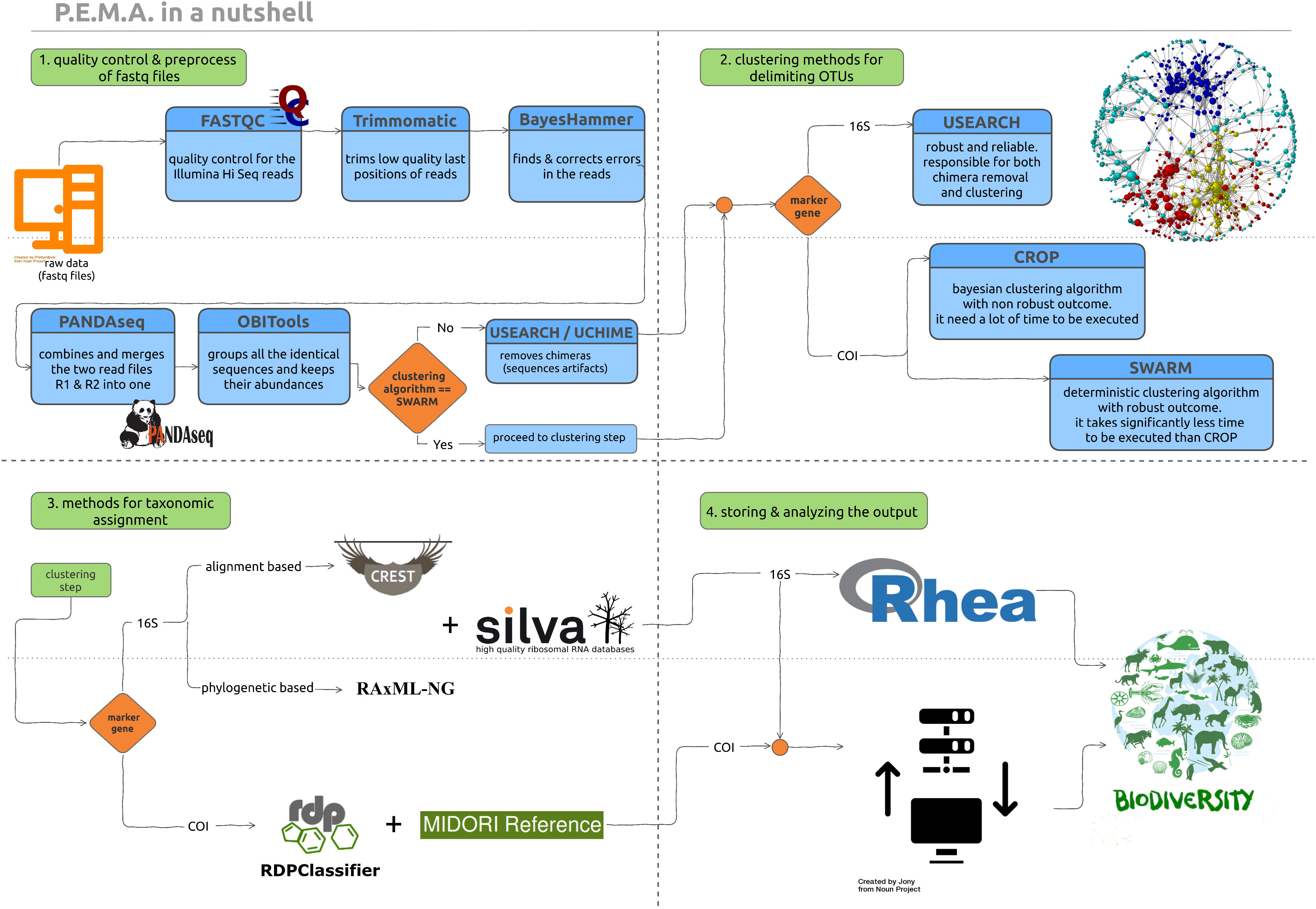
PEMA comprises four parts. The first step (top left) is the quality control and pre-processing of the Illumina sequencing reads. This step is common for both 16S rRNA and COI marker genes. The second step (top right) is the clustering of reads to (M)OTUs. The third step (bottom left) is the taxonomy assignment to the generated (M)OTUs. In the fourth step (bottom right), the results of the metabarcoding analysis are provided to the user and visualized.

### Part 1: Quality control and pre-processing of raw data

Before all else, FastQC [18] is used to obtain an overall read-quality summary. Beyond this visual inspection and to correct the errors are produced by a sequencer, PEMA incorporates a number of tools. Trimmomatic [19] implements a series of trimming steps, namely: either to remove parts of the sequences corresponding to the adapters or the primers, or to trim and crop parts of the reads, or even remove a read completely, when it fails to the quality filtering sequences. BayesHammer [20], an algorithm of the SPAdes assembly toolkit [21], determines specific-position errors, where a particular base has been called incorrectly and revises them. PANDAseq [22] assembles the overlapping paired-end reads and then the ‘obiuniq’ program of OBITools [23] groups all the identical sequences in every sample, keeping a track of their abundances. The VSEARCH package [10] is invoked for the chimera removal.

### Part 2: (M)OTUs clustering

Quality controlled and processed sequences are subsequently clustered into (M)OTUs. For the case of 16S rRNA marker gene, VSEARCH [10] is used. Among the VSEARCH clustering options, PEMA supports *--cluster_size* and *--cluster_unoise.*

For the COI marker gene, two different clustering algorithms are included. First, Swarm v2 [8], a fast and robust algorithm that produces fine-scale MOTUs, free of arbitrary global clustering thresholds and input-order dependency. As the Swarm v2 algorithm is not affected by chimeras (F. Mahé, personal communication), when Swarm v2 is selected, chimera removal occurs after the clustering. This leads to a computational time gain as chimeras are sought among MOTUs instead of unclustered reads.

Second, CROP [9], an unsupervised probabilistic Bayesian clustering algorithm that models the clustering process using Birth-death Markov chain Monte Carlo (MCMC). The CROP clustering algorithm is adjusted by a series of parameters need to be tuned by the user (namely *b, e* and z). These parameters depend on specific dataset properties like the length and the number of reads). PEMA, automatically adjusts *b, e* and *z* by collecting such information and applying the CROP recommended parameter-setting rules [9].

Any singletons occurring among the (M)OTUs after this step are removed.

### Part 3: Taxonomy assignment

Alignment-based taxonomy assignment is supported for both the 16S rRNA and the COI marker gene analyses. In the 16S rRNA marker gene alignment-based case, the LCAClassifier algorithm of the CREST set of resources and tools [11], is used together with the Silva database [12] to assign taxonomy to the OTUs. Two versions of Silva are included in PEMA: 128 (Sept 29, 2016) and 132 (Dec 13, 2017).

For the COI marker gene, PEMA uses the RDPClassifier [13] and the MIDORI reference to assign taxonomy of the MOTUs. The MIDORI dataset [14] contains quality controlled metazoan mitochondrial gene sequences from GenBank [24].

Intended primarily for studies from less explored environments, phylogeny-based assignment is available for 16S rRNA marker gene data. PEMA maps OTUs to a custom reference tree of 1000 Silva-derived consensus sequences (created using RAxML-ng [15] and gappa (phat algorithm) [25], Figure 2A). PaPaRa [26] and EPA-ng [16] combine the OTU clustering output and the reference tree to produce a phylogeny-aware alignment and map the 16S rRNA OTUs to the custom reference tree. Beyond the context of PEMA, users may visualize the output with tree viewers like iTOL [27] (Figure 2B).

**Figure 2:**
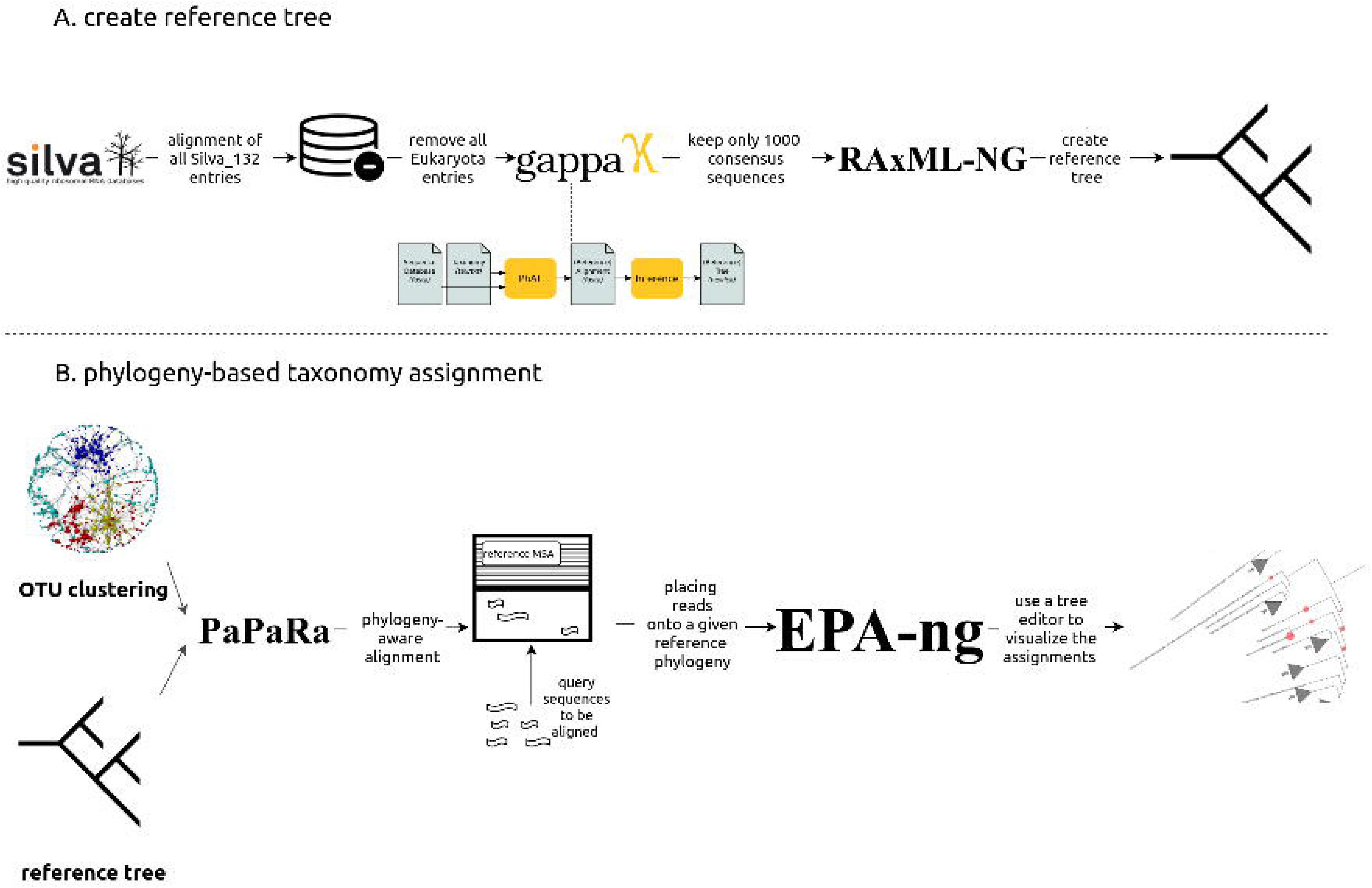
Phylogeny-based taxonomy assignment. A: Building a reference tree for the phylogeny-based taxonomy assignment to 16S rRNA marker gene OTUs: from the latest edition of Silva SSU, all entries referring to Bacteria and Archaea were used and using “art” algorithm, 10000 consensus taxa were kept. B: Using PaPaRa and the OTUs that come up from every analysis, an MSA was made and EPA-ng took over the phylogeny based taxonomy assignment.

### Part 4: (M)OTU and sample abundance tables, biodiversity downstream analysis

In both the 16S rRNA and COI marker gene analysis, an (M)OTU-table is returned by PEMA. For each sample of the analysis, a subfolder containing statistics about the quality of its reads, as well as the taxonomies and their abundances, is also generated.

For the 16S rRNA marker gene case, a downstream analysis of the OTUs including alpha- and beta-diversity analysis, taxonomic composition, statistical comparisons and calculation of correlations between samples is also supported, thanks to the “phyloseq” R package [17]. When this option is selected, then besides phyloseq’s output, a Multiple Sequence Alignment (MSA) and a phylogenetic tree of the OTUs are returned; for the MSA, the MAFFT [28] aligner is invoked while the latter is being built by RAxML-ng [15].

### PEMA container-based installation

An easy way of installing PEMA is via its containers. A dockerized PEMA version is available at https://hub.docker.com/r/hariszaf/pema. Singularity users can *pull* the PEMA image from https://singularity-hub.org/collections/2295. Between the two containers, the Singularity-based one is recommended for HPC environments due to Singularity’s improved security and file accessing properties [29]. For detailed documentation, visit https://github.com/hariszaf/pema.

### PEMA output

All PEMA-related files (i.e. intermediate files, final output, *checkpoint* files and per-analysis-parameters) are grouped in distinct (self-explanatory) subfolders per major PEMA pipeline step. In the last subfolder, the results are further split in folders per sample. This eases further analysis both within the PEMA framework (like partial re-execution for parameter exploration) or beyond.

## Results and discussion

### Evaluation

To evaluate PEMA, two publicly available datasets from published studies were used. For the 16S rRNA marker gene, the dataset reported by study [30] was used while for the COI case, the one of [31] (accession numbers: PRJEB20211 and PRJEB13009 respectively). In both cases, the respective .fastq files were downloaded from ENA-EBI using ‘ENA File Downloader version 1.2’ [32]. For both cases, PEMA was run on the in-house HPC cluster.

### Comparison to existing software

By the means of evaluation, PEMA’s features were compared with those QIIME 2 [2], mothur [1], and LotuS [3]. Table 1 presents a detailed comparison among the four tool features in terms of marker gene support, diversity and phylogeny analysis capability, parameter setting and mode of execution, operation system availability and HPC suitability. As shown, PEMA is equally feature-rich, if not richer in certain feature categories to the other software packages. In particular, PEMA’s support for COI marker gene studies is distinctive; two methods for the taxonomy assignment are supported and PEMA’s easy-parameter setting, step-by-step execution and container distribution render it user and analysis friendly.

**Table 1:**
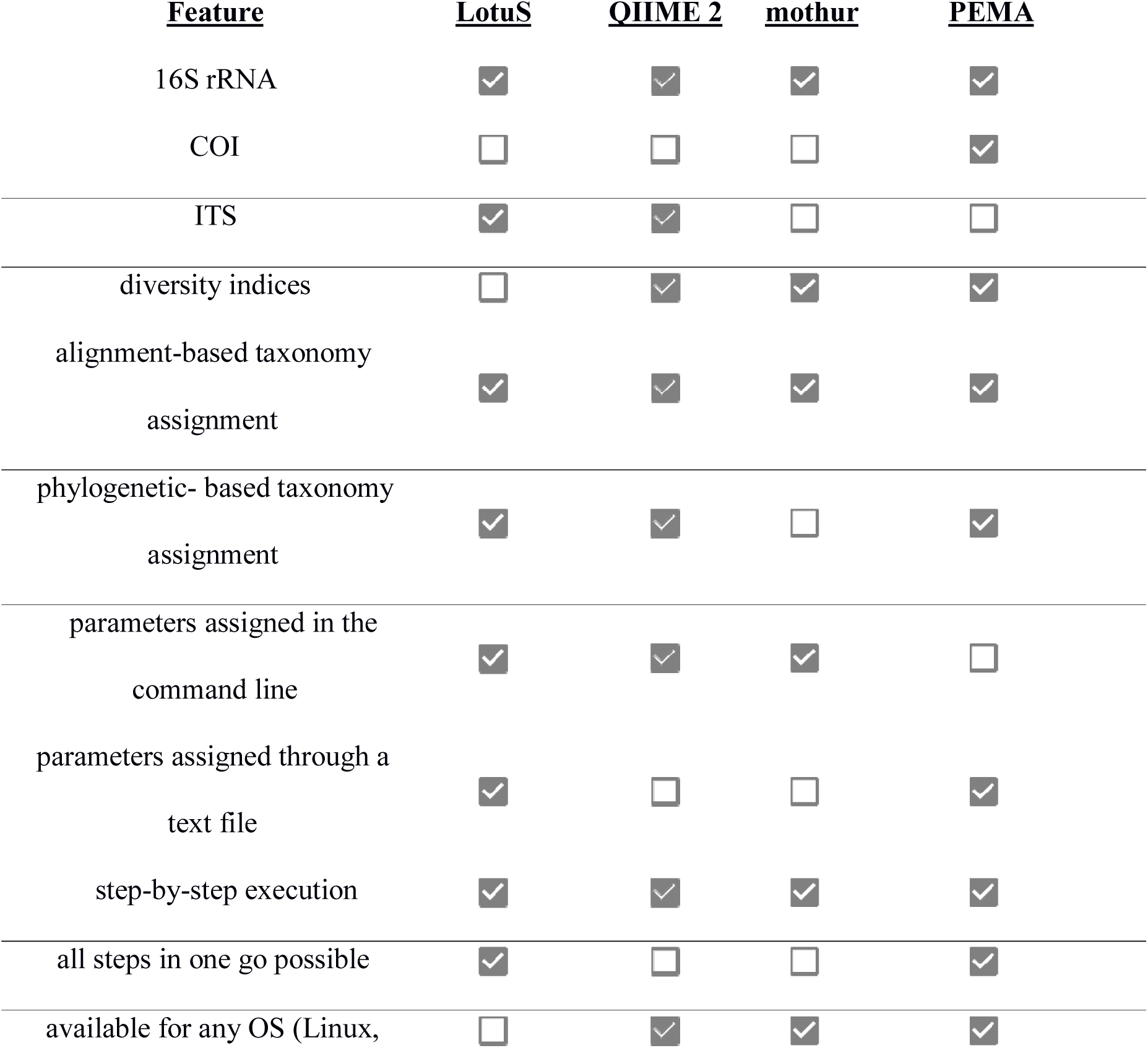
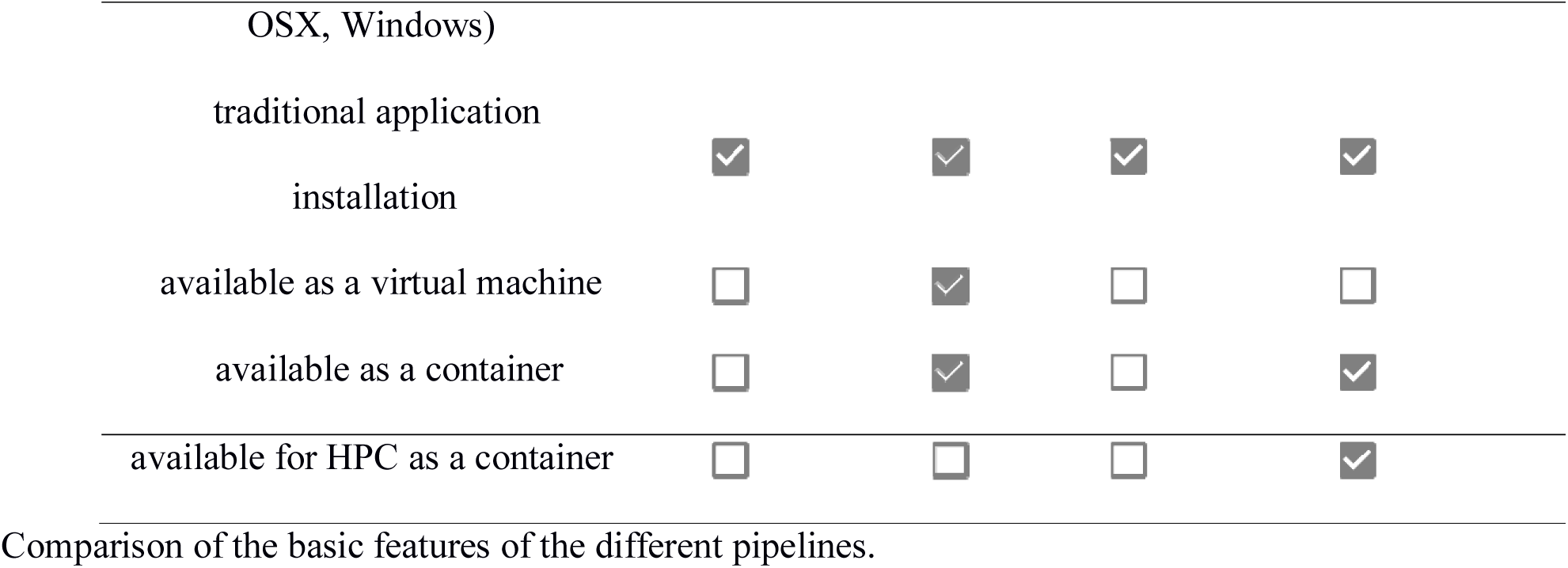
Pipeline comparison.COI marker gene analysis evaluation

The selected study [31] created two COI libraries of different sizes: COIS (235 □bp amplicon size) and COIF (658 □bp amplicon size). The sequencing reads of COIS were selected for PEMA’s evaluation; the COIF sequencing read pairs had no overlap so as to be merged.

Regarding the creation of the MOTU table, [31] used VSEARCH [10] with a clustering at 97% similarity threshold. Afterwards, the BLAST+ (megablast) algorithm [33] was used against a manually created database including all NCBI GenBank COI sequences of length >100 □bp (June 2015) while excluding environmental sequences and higher taxonomic level information [31]. As discussed in the publication, this approach resulted in 138 unique MOTUs out of which 73 were assigned to species level. For PEMA’s evaluation, the chosen clustering algorithm was Swarm v2, using different options for the cluster radius (d) parameter (Table 2); according to [8], this is the most important parameter as it affects the number of MOTUs that are being created. The resulting MOTUs were classified against the MIDORI reference database [14] using RDPClassifier [13]. The results of the processing of the sequences are shown in Additional file 2: Table S1.

**Table 2:** PEMA’s output and executional time.

|  | *d* = 1 | *d* = 2 | *d* = 3 | *d* = 10 | *d* = 13 |
| --- | --- | --- | --- | --- | --- |
| MOTUs after preprocess and clustering steps | 83791 | 59833 | 33227 | 7384 | 4829 |
| MOTUs after chimera removal | 80347 | 57863 | 32539 | 7339 | 4796 |
| Non singletons MOTUs | 6381 | 4947 | 2658 | 1914 | 1634 |
| Assigned species | 62 | 83 | 86 | 86 | 84 |
| Executional time (h) | 02:01:35 | 02:09:49 | 01:51:44 | 02:17:26 | 02:31:15 |

PEMA's output and executional time (using a 20 core node) for different values of Swarm's *d* parameter.

As shown in Table 2, PEMA resulted in 83 species level MOTUs with a cluster radius *(d)* of 2, which is very similar to that of the published study (i.e. 73 species). Although both the clustering algorithm and the taxonomy assignment methods were different between [31] and the present study, the results regarding the number of unique species present in the samples are in agreement to a considerable extent.

The taxonomic assignment of the retrieved MOTUs is shown in Figures 3–4. Certain .fastq files contained very few reads, such as those for sample ERR1308241, and therefore resulted in zero MOTUs upon the completion of PEMA; thus, these samples are not included in Figure 3. It is worth mentioning that four of the 138 MOTUs were found in both the positive control samples of the published study as well as the present study (Table 3). Also, in three cases, PEMA resulted in the same genera as the positive control samples of the published study (Table 3).

**Figure 3:**
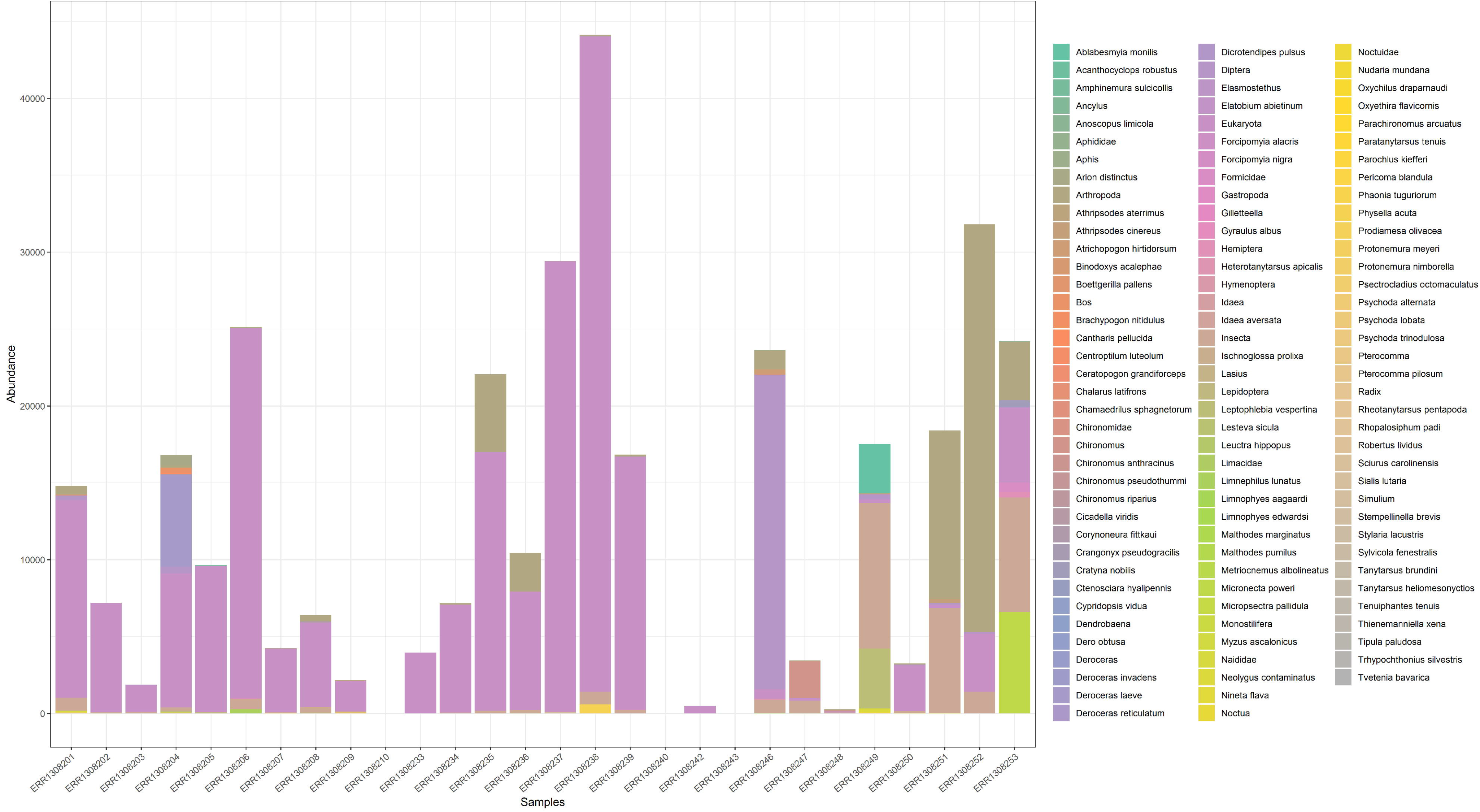
MOTUs bar plot at the lowest possible taxonomic level. Bar plot depicting the taxonomy of the retrieved MOTUs with confidence estimate equal or higher than 0.97 at the lowest possible taxonomic level.

**Figure 4:**
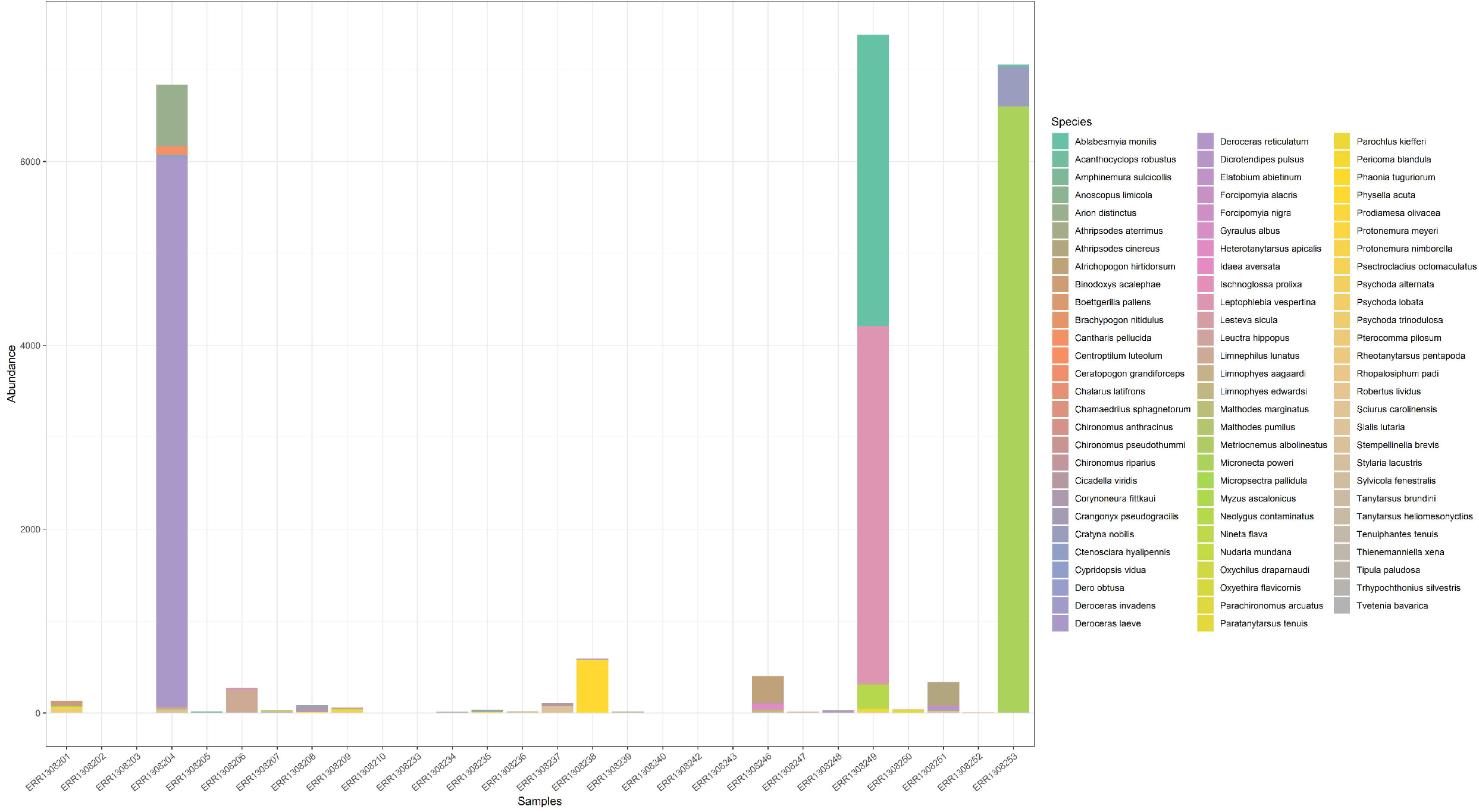
MOTUs bar plot at the species level. Bar plot depicting the taxonomy of the retrieved MOTUs with confidence estimate equal or higher than 0.97 at the species level.

**Table 3:**
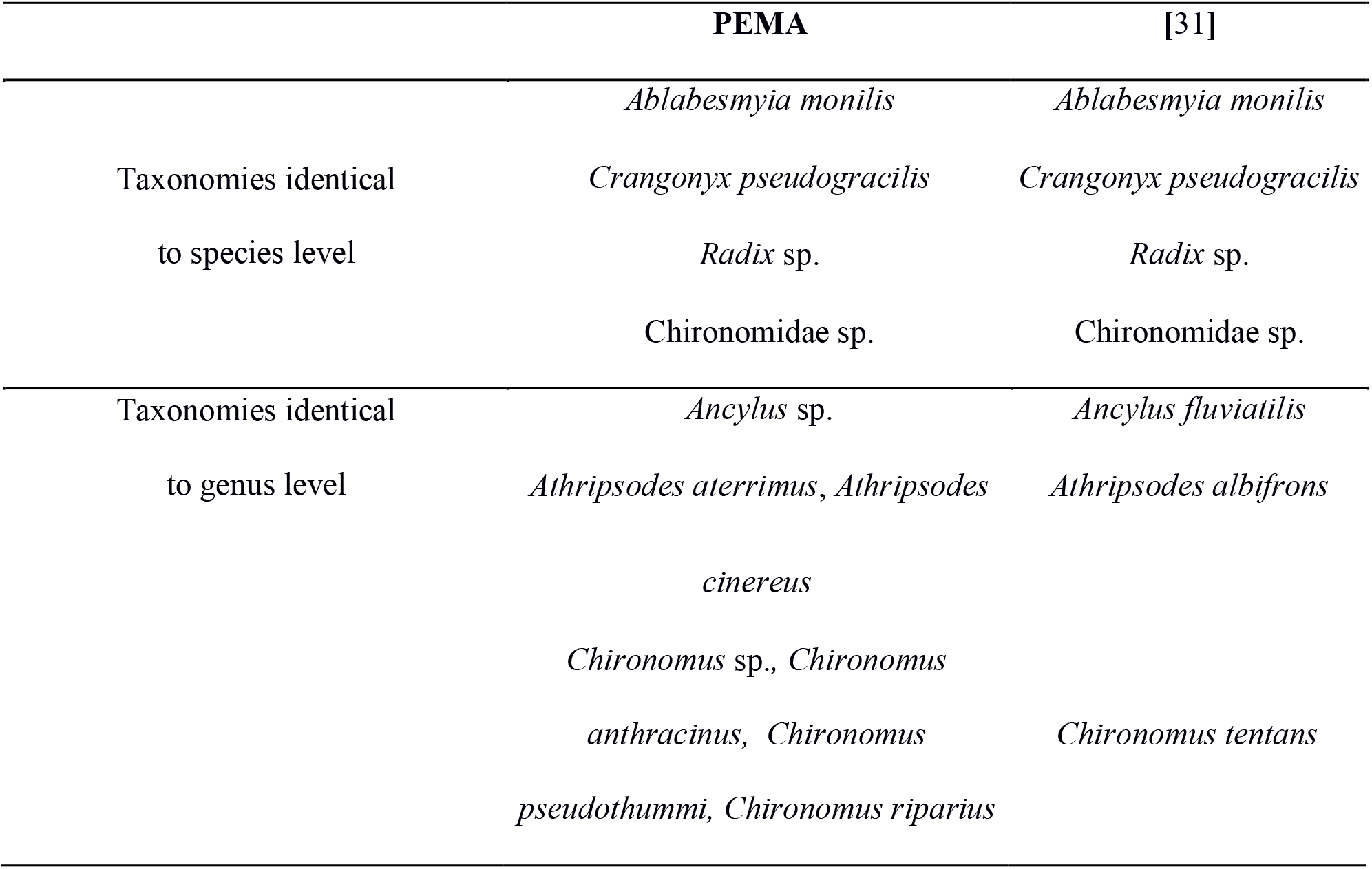
Comparison of the taxonomy of retrieved MOTUs among PEMA and [31].

The computational time required by PEMA for the completion of the analysis is also shown in Table 2. Regardless of the value of the *d* parameter, all analyses were completed in about 2 hours, ie. adequately fast to allow parameter testing and customization.

### 16S rRNA marker gene analysis evaluation

To evaluate PEMA’s performance, a comparative analysis of the [30] dataset with QIIME 2 [2], mothur [1], LotuS [3] and PEMA was conducted.

It is known that the choice of parameters affects the output of each analysis; therefore, it is expected that different user choices might distort the derived outputs. For this reason and for a direct comparison of the pipelines, we have included all the commands and parameters chosen in the framework of this study in the Additional file 1: Supplementary Methods. The results of the processing of the sequences by PEMA are shown in Additional file 3: Table S2. All analyses were conducted on identical Dell M630 nodes (128GB RAM, 20 physical Intel Xeon 2.60GHz cores). LotuS, mothur and QIIME 2 operated in a single thread (core) fashion. PEMA, given the BDS intrinsic parallelization [5], operated with up to the maximum number of node cores (in this case 20).

The execution time and the reported OTU number of each tool are presented in Table 4. LotuS and PEMA resulted in a final number of OTUs comparable to that of [30]. Clearly, due to PEMA’s parallel-execution support, the analysis time can be significantly reduced (~1.5 hours in this case).

**Table 4:**
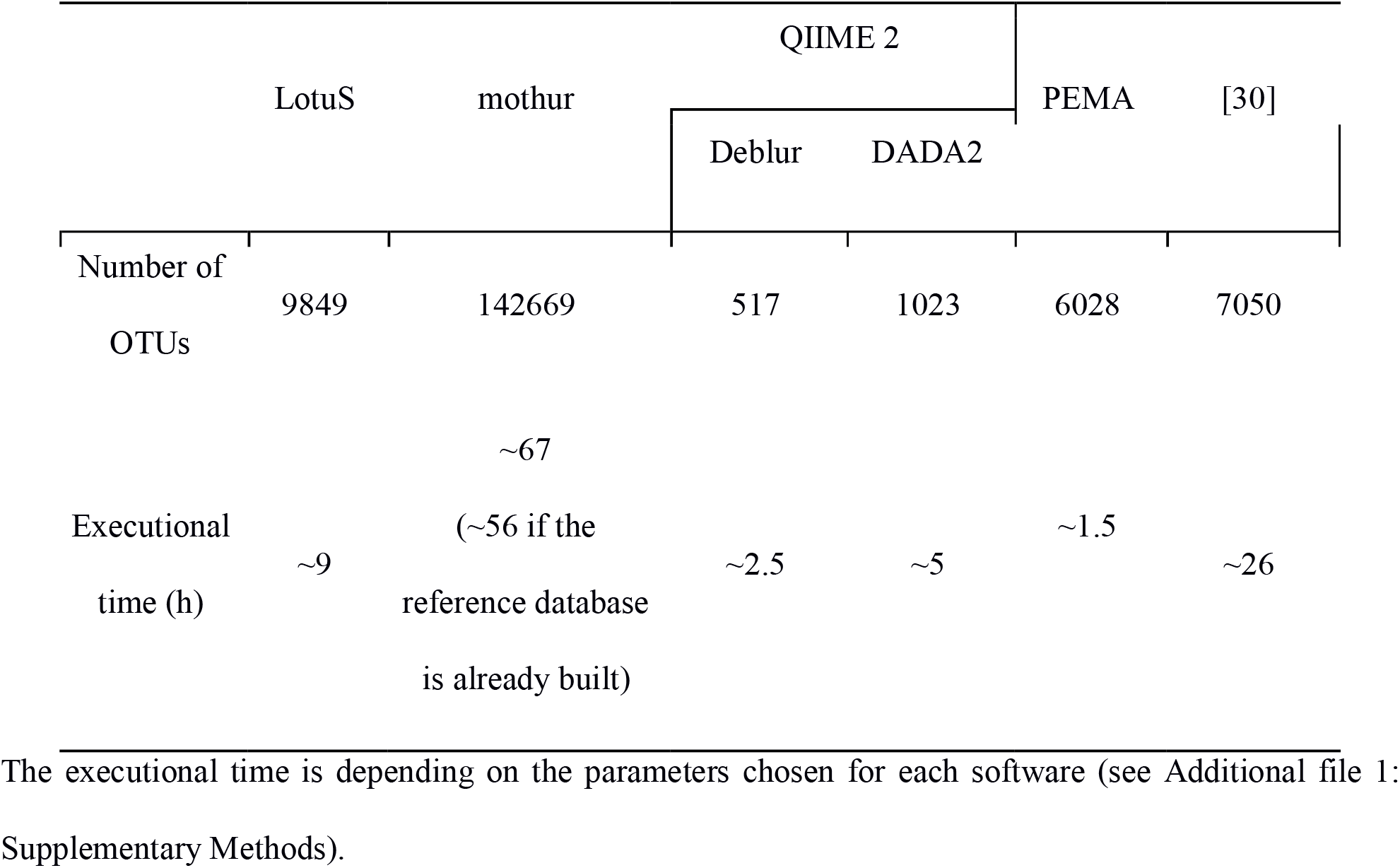
OTU predictions and executional time for the different pipelines.

Due to the non-full overlap of the sequence reads, mothur resulted in an inflated number of OTUs; thus, is was excluded from further analyses. The results of all the pipelines were analyzed with the phyloseq script that is provided with PEMA. The taxonomic assignment of the PEMA retrieved OTUs is shown in Figure 5. The phyla that were found in the samples are similar to the ones that were found in [30]. Although the lowest number of OTUs was found in the marine station (Kal) (Table 5), which is not in accordance with [30], the general trend of the decreasing number of OTUs with the increasing salinity was observed as it was in [30]. Notably, this result was not observed with the other tested pipelines (Table 5). Furthermore, each of the pipelines resulted in a different taxonomic profile (Additional files 4-6: Figure S1-3) with an extreme case of missing the Order of Betaproteobacteriales (Additional files 7-9: Figure S4-6).

**Figure 5:**
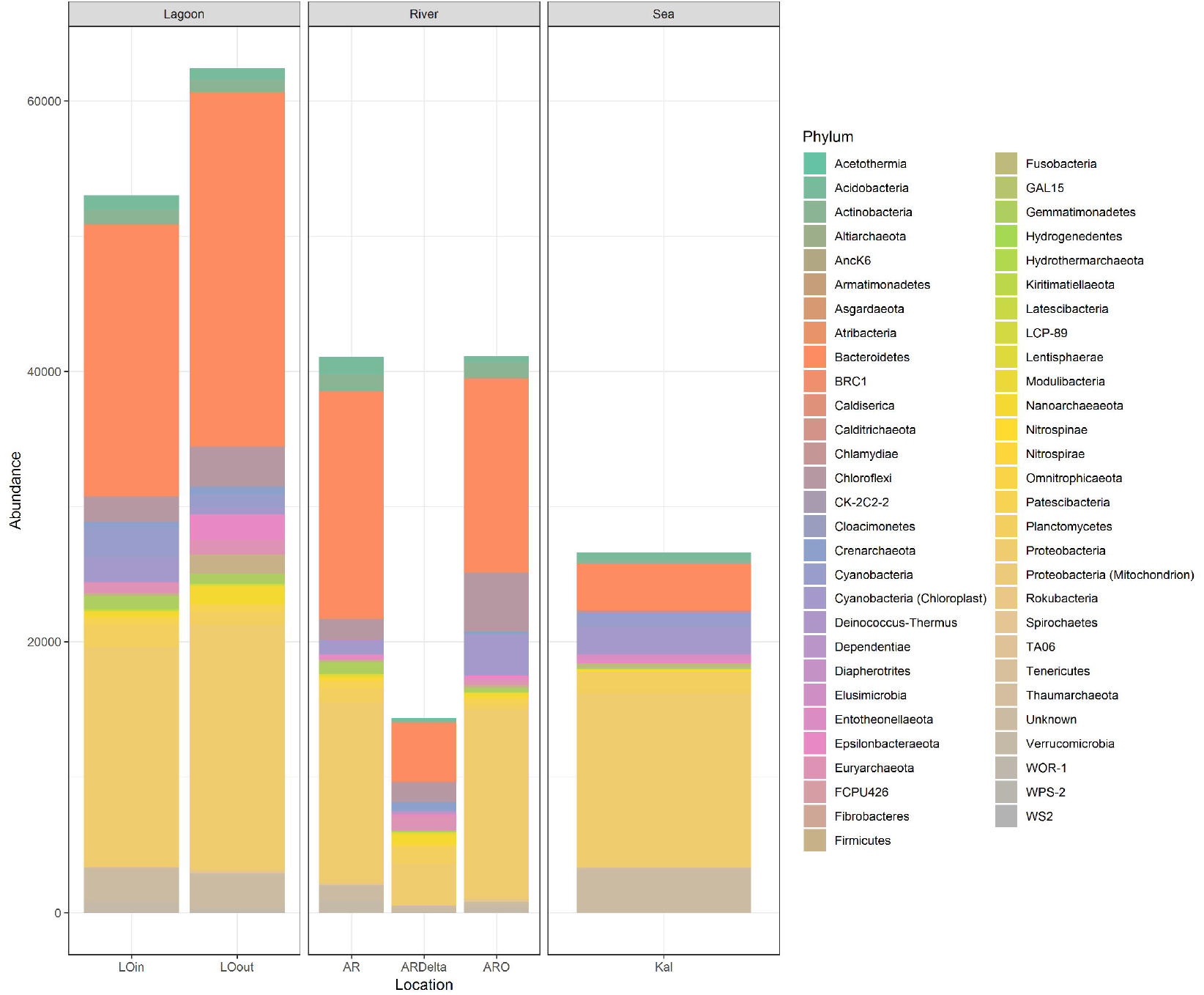
OTUs bar plot at the Phylum level. Bar plot depicting the taxonomy of the retrieved OTUs from PEMA at the Phylum level.

**Table 5:** Diversity indices of the samples.

| Samples | LotuS |  | Deblur |  | DADA2 |  | PEMA |  |
| --- | --- | --- | --- | --- | --- | --- | --- | --- |
|  | OTUs | N | OTUs | N | OTUs | N | OTUs | N |
| L_LOout_A | 2451 | 67640 | 156 | 5878 | 177 | 47954 | 791 | 17888 |
| L_LOout_B | 3432 | 95835 | 180 | 5978 | 248 | 62416 | 1182 | 19571 |
| L_LOout_C | 2987 | 97592 | 180 | 9216 | 221 | 59346 | 1048 | 24947 |
| L_LOinA | 3656 | 85882 | 200 | 6284 | 264 | 62253 | 1176 | 18346 |
| L_LOinB | 2935 | 76545 | 173 | 6357 | 194 | 46191 | 954 | 18750 |
| L_LOinC | 3149 | 71222 | 183 | 5849 | 219 | 49890 | 1012 | 15940 |
| S_KalA | 2467 | 59039 | 82 | 2991 | 173 | 46418 | 668 | 10086 |
| S_KalB | 2940 | 57091 | 89 | 2107 | 192 | 49715 | 802 | 7702 |
| S_KalC | 2898 | 61191 | 101 | 2934 | 214 | 51321 | 769 | 8840 |
| R_ARDelta_A | 3079 | 53780 | 97 | 1341 | 259 | 53237 | 737 | 4386 |
| R_ARDelta_B | 3671 | 63857 | 117 | 1893 | 271 | 62477 | 836 | 5716 |
| R_ARDelta_C | 3323 | 50640 | 100 | 1389 | 230 | 48650 | 703 | 4247 |
| R_AR_A | 3084 | 58311 | 152 | 3000 | 269 | 52689 | 1058 | 10208 |
| R_AR_B | 4011 | 84370 | 199 | 3931 | 304 | 81589 | 1546 | 14058 |
| R_AR_C | 3674 | 97462 | 196 | 5290 | 343 | 83000 | 1501 | 16819 |
| R_ARO_A | 3079 | 63781 | 123 | 1993 | 324 | 60752 | 1000 | 7381 |
| R_ARO_B | 3178 | 92008 | 183 | 5352 | 359 | 64824 | 1307 | 17052 |
| R_ARO_C | 3248 | 89992 | 186 | 5540 | 307 | 61082 | 1255 | 16730 |

OTUs: total number of OTUs. N: total microbial relative abundance values. All sample libraries that start with “L” correspond to samples collected from lagoons, with “S” from the sea while “R” stands for the riverine samples. AR: Arachthos. ARO: Arachthos Neochori. ARDelta: Arachthos Delta. LOin: Logarou station inside the lagoon. LOout: Logarou station in the channel connecting the lagoon to the gulf. Kal: Kalamitsi. A, B, C replicate samples.

Moreover, when the PERMANOVA analysis was run for the results of PEMA, LotuS and DADA2, it was clear that the microbial community composition was significantly different in each of the three sampled habitats (i.e. River, Lagoon, Sea) (PERMANOVA: F.Model = 7.0718, p < 0.001; F.Model = 6.5901, p < 0.001, F.Model = 2.2484, p < 0.05, respectively), which is in accordance with [30]. However, this was not the case with Deblur (PERMANOVA: p > 0.05).

Overall, PEMA’s output is in accordance with [30] and if all the different outputs are taken into consideration, it can be concluded that it performed better than the other tested pipelines in capturing the microbial community diversity and composition of the samples.

### Beyond environmental ecology, on-going and future work

PEMA is mainly intended to support eDNA metabarcoding analysis and be directly applicable to next-generation biodiversity/ecological assessment studies. Given that community composition analysis may also serve additional research fields, eg. microbial pathology, the potential impact of such pipelines is expected to be much higher. On-going PEMA work focuses on serving a wide scientific audience and on making it applicable to more types of studies. The easy set up and execution of PEMA, allows users to work closely with national and European HPC/e-infrastructures (e.g. ELIXIR Greece [34], LifeWatch ERIC[35], EMBRC ERIC[36]). The aim of this effort is to outreach their communities and address both ongoing as well as future analysis needs.

In a mid-to meso-term perspective, pipeline development work will support the analysis of the ITS and 18S rRNA marker gene studies too. When operational, a holistic biodiversity assessment approach would be possible through PEMA and eDNA; the analysis of the most commonly used marker genes (16S rRNA, ITS, COI/18S rRNA) for each of the greatest taxonomic groups (Bacteria, Fungi, Eukaryotes) would be implemented by the same pipeline. Finally, it is our intention to allow *ad hoc* and in-house databases to be used as reference for the (M)OTU taxonomy assignment.

## Conclusions

PEMA is an accurate, execution friendly and fast pipeline for the metabarcoding analysis of the 16S rRNA and COI marker genes. It provides a per-sample analysis output, different taxonomy assignment methods and graphics-based biodiversity/ecological analysis. This way, in addition to (M)OTU calling, it provides users with both an informative study overview and detailed result snapshots.

PEMA’s user friendliness derives from the easy and with minimal number of installation and execution commands. In addition, PEMA’s strategic choice of a single parameter file, implementation programming language, and multiple container-type distribution, grant it with speed (running in parallel), on-demand partial pipeline enactment, and provision for HPC-system-based sharing.

All the aforementioned features, render PEMA attractive for biodiversity/ecological assessment analyses. Applications may mainly concern environmental ecology with possible extensions to fields like microbial pathology and gut microbiome, inline with modern research needs, from low volume to big data.

## Availability of supporting source code and requirements

Project name: PEMA

Project home page: https://github.com/hariszaf/pema

Archived version: see project home page (github repository)

Operating system(s): Platform independent

Programming language: BigDataScript

Other requirements: Singularity (in case of HPC usage)

License: GNU GPLv3 (for 3rd party components separate licenses apply)

Any restrictions to use by non-academics: licence needed

## Availability of supporting data

The sequence data that support the findings of this study are available in European Nucleotide Archive (ENA) with the study accession numbers PRJEB20211 (http://www.ebi.ac.uk/ena/data/view/PRJEB20211) and PRJEB13009 (https://www.ebi.ac.uk/ena/data/view/PRJEB13009).

## Declarations

## List of abbreviations

BDS: BigDataScript
COI: Cytochrome Oxidase Subunit 1
eDNA: Environmental DNA
MOTU: Molecular Operational Taxonomic Unit (species equivalent for Eukaryotes)
HPC: High Performance Computing
MCMC: Markov chain Monte Carlo
MSA: Multiple Sequence Alignment
OTU: Operational Taxonomic Unit (species equivalent for prokaryotes)
PEMA: a Pipeline for Environmental DNA Metabarcoding Analysis
SSU: Small Subunit

## Ethics approval and consent to participate

Not applicable

## Consent for publication

Not applicable

## Authors’ contributions

HZ conceived and designed the pipeline, performed its containerization, analyzed and interpreted the data, wrote the paper, prepared figures and/or tables, reviewed drafts of the paper. HQV offered support in the HPC preparation and setup and in 3rd party component usage. KV and CA conceived the idea and reviewed drafts of the paper. PT conceived the idea, proposed the usage of the programming language and reviewed drafts of the paper. CP conceived the idea, prepared figures and/or table and reviewed drafts of the paper. AP offered support in HPC and in 3rd party components. EP conceived the idea, assisted with programming and setup and reviewed drafts of the paper. All authors read and approved the final manuscript.

## Acknowledgements

The authors would like to thank: a. the Information technology (IT) group of HCMR and especially Mr Stelios Ninidakis, Mr Georgios Tsamis and Mr Dimitris Sidirokastritis for their help and support during cluster maintenance and installation of third party software. b. Dr. Christos A. Christakis (ORCID iD: 0000-0002-7075-0996) for his valuable feedback on ecological analysis usefulness aspects.

## Additional files

**Additional file 1: Supplementary Methods:** Description of tools invoked by PEMA and their licences. Description of the commands, along with their parameters, used to run PEMA, mothur, LotuS and QIIME 2.

**Additional file 2: Table S1:** Number of sequences after each pre-processing step for the case of COI, dataset from [31].

**Additional file 3: Table S2:** Number of sequences after each pre-processing step for the case of 16S rRNA gene.

**Additional file 4: Figure S1:** Bar plot depicting the taxonomy of the retrieved OTUs from LotuS at the Phylum level.

**Additional file 5: Figure S2:** Bar plot depicting the taxonomy of the retrieved OTUs from QIIME 2 using Deblur at the Phylum level.

**Additional file 6: Figure S3:** Bar plot depicting the taxonomy of the retrieved OTUs from QIIME 2 using DADA2 at the Phylum level.

**Additional file 7: Figure S4:** Bar plot depicting the taxonomy of the retrieved OTUs from LotuS at the class of Betaproteobacteriales.

**Additional file 8: Figure S5:** Bar plot depicting the taxonomy of the retrieved OTUs from QIIME 2 using Deblur at the class of Betaproteobacteriales.

**Additional file 9: Figure S6:** Bar plot depicting the taxonomy of the retrieved OTUs from PEMA at the class of Betaproteobacteriales.

## References

[1] Schloss PD, Westcott SL, Ryabin T, Hall JR, Hartmann M, Hollister EB, et al. Introducing mothur: open-source, platform-independent, community-supported software for describing and comparing microbial communities. Appl. Environ. Microbiol. 2009; 75:7537–41.

[2] Bolyen E, Rideout JR, Dillon MR, Bokulich NA, Abnet C, Al-Ghalith GA, et al. QIIME 2: Reproducible, interactive, scalable, and extensible microbiome data science. PeerJ Preprints. 2018; 6:e27295v2.

[3] Hildebrand F, Tadeo R, Voigt AY, Bork P, Raes J. LotuS: an efficient and user-friendly OTU processing pipeline. Microbiome. 2014; 2:30.

[4] European Strategy Forum on Research Infrastructures Innovation Working Group. Innovation-oriented cooperation of Research Infrastructures. Vol.3. ESFRI Scripta. 2018.

[5] Cingolani P, Sladek R, Blanchette M. BigDataScript: a scripting language for data pipelines. Bioinformatics. 2014; 31:10–16.

[6] Rad BB, Bhatti HJ, Ahmadi M. An introduction to docker and analysis of its performance. International Journal of Computer Science and Network Security (IJCSNS). 2017; 17:228.

[7] Kurtzer GM, Sochat V, Bauer MW. Singularity: Scientific containers for mobility of compute. PloS one. 2017; 12:e0177459.

[8] Mahé F, Rognes T, Quince C, de Vargas C, Dunthorn M. Swarm v2: highly-scalable and high-resolution amplicon clustering. PeerJ. 2015; 3:e1420.

[9] Hao X, Jiang R, Chen T. Clustering 16S rRNA for OTU prediction: a method of unsupervised bayesian clustering. Bioinformatics. 2011; 27:611–8.

[10] Rognes T, Flouri T, Nichols B, Quince C, Mahé F. VSEARCH: a versatile open source tool for metagenomics. PeerJ. 2016; 4:e2584.

[11] Lanzén A, Jørgensen SL, Huson DH, Gorfer M, Grindhaug SH, Jonassen I, et al. CREST-classification resources for environmental sequence tags. PloS one. 2012; 7:e49334.

[12] Quast C, Pruesse E, Yilmaz P, Gerken J, Schweer T, Yarza P, Peplies J, Glöckner FO. The SILVA ribosomal RNA gene database project: improved data processing and web-based tools. Nucl. Acids Res. 2013; 41:D590–6.

[13] Wang Q, Garrity GM, Tiedje JM, Cole JR. Naive Bayesian classifier for rapid assignment of rRNA sequences into the new bacterial taxonomy. Appl. Environ. Microbiol. 2007; 73:5261–7.

[14] Machida RJ, Leray M, Ho SL, Knowlton N. Metazoan mitochondrial gene sequence reference datasets for taxonomic assignment of environmental samples. Scientific data. 2017; 4:170027.

[15] Kozlov AM, Darriba D, Flouri T, Morel B, Stamatakis A. RAxML-NG: a fast, scalable and user-friendly tool for maximum likelihood phylogenetic inference. Bioinformatics. 2019; btz305.

[16] Barbera P, Kozlov AM, Czech L, Morel B, Darriba D, Flouri T, Stamatakis A. EPA-ng: massively parallel evolutionary placement of genetic sequences. Systematic biology. 2018; 68:365–9.

[17] McMurdie JP, Holmes S. phyloseq: an R package for reproducible interactive analysis and graphics of microbiome census data. PloS one. 2013; 8:e61217.

[18] Andrews S. FastQC. http://www.bioinformatics.babraham.ac.uk/projects/fastqc/. Accessed 08 July 2019.

[19] Bolger AM, Lohse M, Usadel B. Trimmomatic: a flexible trimmer for illumina sequence data. Bioinformatics. 2014; 30:2114–20.

[20] Nikolenko SI, Korobeynikov AI, Alekseyev MA. Bayeshammer: Bayesian clustering for error correction in single-cell sequencing. BMC genomics. 2013; S7.

[21] Bankevich A, Nurk S, Antipov D, Gurevich AA, Dvorkin M, Kulikov AS, et al. Spades: a new genome assembly algorithm and its applications to single-cell sequencing. Journal of computational biology. 2012; 19:455–77.

[22] Masella AP, Bartram AK, Truszkowski JM, Brown DG, Neufeld JD. PANDAseq: paired-end assembler for illumina sequences. BMC bioinformatics. 2012; 13:31.

[23] Boyer F, Mercier C, Bonin A, Le Bras Y, Taberlet P, Coissac E. OBITools: a unix-inspired software package for dna metabarcoding. Molecular ecology resources. 2016; 16:176–82.

[24] Benson DA, Cavanaugh M, Clark K, Karsch-Mizrachi I, Ostell J, Pruitt KD, Sayers EW. GenBank. Nucleic acids research 2018; 46:D41–47.

[25] Czech L, Barbera P, Stamatakis A. Methods for automatic reference trees and multilevel phylogenetic placement. Bioinformatics. 2018; 35:1151–8.

[26] Berger SA, Stamatakis A. PaPaRa 2.0: a vectorized algorithm for probabilistic phylogeny-aware alignment extension. Heidelberg Institute for Theoretical Studies. 2012.

[27] Letunic I, Bork P. Interactive tree of life (itol): an online tool for phylogenetic tree display and annotation. Bioinformatics. 2006; 23:127–8.

[28] Katoh K, Misawa K, Kuma KI, Miyata T. Mafft: a novel method for rapid multiple sequence alignment based on fast fourier transform. Nucleic acids research. 2002; 30:3059–66.

[29] Chavez J. Singularity: a “Docker” for HPC environments. https://dev.to/grokcode/singularity--a-docker-for-hpc-environments-i6p. Accessed 08 Jul 2019.

[30] Pavloudi C, Kristoffersen JB, Oulas A, De Troch M, Arvanitidis C. Sediment microbial taxonomic and functional diversity in a natural salinity gradient challenge Remane’s “species minimum” concept. PeerJ. 2017; 5:e3687.

[31] Bista I, Carvalho GR, Walsh K, Seymour M, Hajibabaei M, Lallias D, et al. Annual time-series analysis of aqueous edna reveals ecologically relevant dynamics of lake ecosystem biodiversity. Nature communications. 2017; 8:14087.

[32] Harrison PW, Alako B, Amid C, Cerdeño-Tárraga A, Cleland I, Holt S, et al. The European Nucleotide Archive in 2018. Nucleic acids research. 2018; 47:D84–8.

[33] Camacho C, Coulouris G, Avagyan V, Ma N, Papadopoulos J, Bealer K, Madden TL BLAST+: architecture and applications. BMC bioinformatics. 2009; 10:421.

[34] ELIXIR-GR. https://www.elixir-greece.org/ Accessed 08 July 2019.

[35] LifeWatch-ERIC. https://www.lifewatch.eu/ Accessed 08 July 2019.

[36] EMBRC. http://www.embrc.eu/ Accessed 08 July 2019.

