## Supplementary Methods for "PEMA: from the raw .fastq files of 16S rRNA and COI marker genes to the (M)OTU-table, a thorough metabarcoding analysis"

Each of the tools PEMA uses has some special characteristics that answer to the amplicon analysis’ demands in a better way, compared to others. In this supplementary file, some of these characteristics are recorded.

***FastQC***

After a dataset has been analysed using PEMA, samples whose .fastq files are of low quality can be excluded from subsequent PEMA runs of this particular dataset. However, removing files completely does not secure that all the remaining reads are of satisfactory quality, as either smaller regions of a read or specific-position errors might have occurred during the sequencing. The next tools in PEMA address these issues.

***Trimmomatic***

As PEMA deals with amplicons, the handling of technical sequences erroneously produced during the sequencing is crucial. Amplicons in particular could cause significant alterations due to their common presence in the end of the reads. Trimmomatic addresses the need of removing such sequences.

When run in ‘Simple mode’, Trimmomatic aims to find approximate matches between the reads and the technical sequences supplied by the user. This way, technical sequences can be identified in any location or orientation within the reads. A considerable overlap between the read and a technical sequence is required to prevent false-positive findings. However, in some cases, the sequenced DNA fragment is shorter than the read length. For these cases, Trimmomatic’s ‘Palindrome mode’ uses both reads and, given the latter are reverse complements, is able to locate short partial adapter sequences; sequences that would otherwise remain undetected as they miss the minimum required overlap.

Trimmomatic, in the aforementioned ways, detects and removes regions of low sequencing quality. However, it is not able to correct a wrong read base. For this task the next algorithm was recruited.

***BayesHammer***

BayesHammer algorithm tries to

1) find read k-mers whose sequencing quality is above a predefined threshold (“solid k-mers”), and

2) select the sequencing reads that can be rebuilt based on the aforementioned “solid k-mers”.

Thus, after finding the “solid k-mers”, BayesHammer can decide whether a base is sequenced properly or not.

***PANDAseq***

PANDAseq attempts to correct possible errors using a probabilistic approach based on the overlap data from the paired-end reads. Further, it uses intrinsic properties of Illumina sequencing, including the low probability of gap inclusion. Finally, it also takes into account the primers that have been used.

PANDAseq supports more than one merging algorithms. ‘Pear’ and ‘simple\_bayesian’ are the ones used most commonly. In PEMA the default merging algorithm is set in ‘simple\_bayesian’; shifting to another PANDAseq-supported algorithm is possible for users that would like to optimize this step via experimentation.

***OBITools***

OBITools is a set of python programs developed to simplify the manipulation of sequence files, especially in the context of NGS-based DNA metabarcoding.

PEMA uses the ‘obiuniq’ program of OBITools with the PANDAseq merged files. A set of fastq files is returned. In the header of each entry, except from the sample name and the sequence id there is also the number of copies of this sequence that were found in this sample.

***VSEARCH***

VSEARCH is an open source and free of charge tool that is meant to be used as an alternative to the USEARCH tool; as on the one hand USEARCH source code is not publicly available and on the other, it is not free distributed.

VSEARCH achieves the processing and preparing of sequence data that arrive from a series of different fields like metagenomics, genomics and population genomics. Among other tasks, VSEARCH supports chimera removal, clustering and dereplication of the sequencing reads. VSEARCH also supports multithreading and as a result takes advantage of this parallelism to perform accurate results at high speed.

***Swarm v2***

The way Swarm builds OTUs should not be affected by chimeras, according to the developers of the algorithm. Chimeras will very likely form independent OTUs. Hence, PEMA checks for chimeras after clustering when Swarm is used. Swarm creates a fasta file containing ΜOTU representatives and with this file PEMA runs a chimera check, using again VSEARCH algorithm. With the output of ‘uchime3_denovo’, PEMA builds an OTU-table without taxonomies.

In the “.stats” file that Swarm produces, the number of unique amplicons in the MOTU, the total abundance of amplicons in it, the identifier of the initial seed, the abundance of that initial seed and the number of amplicons with abundance 1 stand for the first 6 columns of “.stats” file, respectively. The last two columns represent the maximum number of iterations before the MOTU reached its natural limit and the cumulative number of steps along the path joining the seed and the furthermost amplicon in the MOTU. Furthermore, in case that parameter d equals to 1, all first five columns are modified, as a second clustering pass is performed to reduce the number of small MOTUs, but not the last two.

***CREST & SILVA***

The ‘LCAClassifier’ algorithm is used for the classification of sequences aligned to the reference databases provided. CREST can handle 4 databases: SILVA, Greengenes, Unite and amoA. Moreover, it allows the user to build his/her own database.

***BigDataScript programming language (BDS)***

PEMA is a very fast pipeline for a metabarcoding analysis. This is mainly due to the BDS programming language that allows absolute serialization and lazy processing. This way, even in the case that an error takes place, PEMA is able to restart from the last checkpoint that was created, before the error occurred. In addition, when PEMA has to perform a “for” loop, then BDS splits that task in all cores available. Then, thanks to the “wait” command that BDS includes, it just stops until everything needed for the next step of the pipeline is completed. However, even if BDS has a lot of advantages, some of the tools included in BDS cannot exploit them. That is why, even if in some steps there are nodes available, PEMA does not use them.

After a failure of a task, the program can be re-executed from that specific point and there is no need to re-run it from scratch. This is also possible in case a pipeline has more than one possible arguments in a specific step. Re-executing the pipeline from a specific step, after changing a specific parameter, is possible with BDS checkpoints. This is quite convenient for a pipeline like PEMA as after the completion of an analysis, if the user wishes for a single parameter of a single step to be changed, PEMA can be executed only from the point that this parameter appears and not earlier. For example, even if P.E.M.A has been executed with Swarm as clustering algorithm, BDS checkpoints allow the user to rerun PEMA with CROP as clustering algorithm, only from the clustering step and forward.

**Licenses**

- BDS	Apache License 2 (<http://www.apache.org/licenses/LICENSE-2.0>)
- Trimmomatic, PANDAseq,	GNU GPLv3 (<https://www.gnu.org/licenses/gpl-3.0.en.html>)
  PaPaRa, VSEARCH,
  CREST, Fastqc, RAxML-ng,
  Crop, EPA-ng
- Phyloseq, vegan, Swarm v2	AGPLv3 (<https://opensource.org/licenses/AGPL-3.0>)
- Blastn (NCBI BLAST)	Public Domain Notice ([https://www.ncbi.nlm.nih.gov/IEB/ToolBox
  /CPP_DOC/lxr/source/scripts/projects/blast/LICENSE](https://www.ncbi.nlm.nih.gov/IEB/ToolBox/CPP_DOC/lxr/source/scripts/projects/blast/LICENSE))
- Spades, RDPTools	GNU GPLv2 (<https://opensource.org/licenses/GPL-2.0> )
- OBITools	cecill ([http://www.cecill.info/licences/Licence_CeCILL_V2.1
  -en.html](http://www.cecill.info/licences/Licence_CeCILL_V2.1-en.html))
- Mafft	BSD license (<https://mafft.cbrc.jp/alignment/software/license.txt>)

### **Comparison of pipelines**

For reference, we include the commands used to execute QIIME 2, mothur, LotuS and PEMA on the chosen dataset. The official QIIME 2 tutorial (<https://docs.qiime2.org/2019.4/tutorials/moving-pictures/>), mothur SOP (<https://www.mothur.org/wiki/MiSeq_SOP>) and LotuS tutorial (<http://psbweb05.psb.ugent.be/lotus/documentation.html#Tutorial>) were followed.

***QIIME 2 commands***

In order to import our data we ran:

```
qiime tools import \
  --type 'SampleData[PairedEndSequencesWithQuality]' \
  --input-path pe-64-manifest \
  --output-path paired-end-demux.qza \
  --input-format PairedEndFastqManifestPhred64V2
```

Then, the two different approaches that QIIME 2 supports were both performed.

Hence, for the DADA2 approach, the below commands were performed:

```
qiime dada2 denoise-single \
  --i-demultiplexed-seqs paired-end-demux.qza \
  --p-trim-left 0 \
  --p-trunc-len 120 \
  --o-representative-sequences rep-seqs-dada2.qza \
  --o-table table-dada2.qza \
  --o-denoising-stats stats-dada2.qza

qiime metadata tabulate \
  --m-input-file stats-dada2.qza \
  --o-visualization stats-dada2.qzv
mv rep-seqs-dada2.qza rep-seqs.qza
mv table-dada2.qza table.qza
```

While for the Deblur approach, the following commands were executed instead:

```
qiime quality-filter q-score \
 --i-demux paired-end-demux.qza \
 --o-filtered-sequences demux-filtered.qza \
 --o-filter-stats demux-filter-stats.qza
qiime deblur denoise-16S \
  --i-demultiplexed-seqs demux-filtered.qza \
  --p-trim-length 120 \
  --o-representative-sequences rep-seqs-deblur.qza \
  --o-table table-deblur.qza \
  --p-sample-stats \
  --o-stats deblur-stats.qza
qiime metadata tabulate \
  --m-input-file demux-filter-stats.qza \
  --o-visualization demux-filter-stats.qzv
qiime deblur visualize-stats \
  --i-deblur-stats deblur-stats.qza \
  --o-visualization deblur-stats.qzv
mv rep-seqs-deblur.qza rep-seqs.qza
mv table-deblur.qza table.qza
```

After that, for both cases the succeeding commands were performed:

In order to get feature table and feature data summaries:

```
qiime feature-table summarize \
  --i-table table.qza \
  --o-visualization table.qzv \
  --m-sample-metadata-file sample-metadata.tsv
qiime feature-table tabulate-seqs \
  --i-data rep-seqs.qza \
  --o-visualization rep-seqs.qzv
```

And for the taxonomy assignment of the OTUs found:

```
qiime feature-classifier classify-sklearn \
  --i-classifier gg-13-8-99-515-806-nb-classifier.qza \
  --i-reads rep-seqs.qza \
  --o-classification taxonomy.qza

qiime metadata tabulate \
  --m-input-file taxonomy.qza \
  --o-visualization taxonomy.qzv
```

***mothur commands***

The mothur sbatch file was set as such:

#This is a batch file for running the equivalent of mothur

make.file(inputdir=/home1/haris/MyData_mothur/, type=fastq, prefix=mapping)

make.contigs(file=mapping.files, processors=8)

summary.seqs(fasta=mapping.trim.contigs.fasta)

screen.seqs(fasta=mapping.trim.contigs.fasta, group=mapping.contigs.groups, summary=mapping.trim.contigs.summary, maxambig=0, minlength=100, maxlength=600, maxhomop=8)

unique.seqs(fasta=mapping.trim.contigs.good.fasta)

count.seqs(name=mapping.trim.contigs.good.names, group=mapping.contigs.good.groups)

summary.seqs(count=mapping.trim.contigs.good.count_table)

pcr.seqs(fasta=silva.nr_v132.align, start=1044, end=43116, keepdots=T, processors=8)

rename.file(input=silva.nr_v132.pcr.align, new=silva.fasta)

summary.seqs(fasta=silva.fasta)

align.seqs(fasta=mapping.trim.contigs.good.unique.fasta, reference=silva.fasta)

summary.seqs(fasta=mapping.trim.contigs.good.unique.align, count=mapping.trim.contigs.good.count_table)

screen.seqs(fasta=mapping.trim.contigs.good.unique.align, count=mapping.trim.contigs.good.count_table, summary=mapping.trim.contigs.good.unique.summary, start=6388, end=25316, maxhomop=8)

summary.seqs(fasta=mapping.trim.contigs.good.unique.good.align, count=mapping.trim.contigs.good.good.count_table)

filter.seqs(fasta=mapping.trim.contigs.good.unique.good.align, vertical=T, trump=.)

unique.seqs(fasta=mapping.trim.contigs.good.unique.good.filter.fasta, count=mapping.trim.contigs.good.good.count_table)

pre.cluster(fasta=mapping.trim.contigs.good.unique.good.filter.unique.fasta, count=mapping.trim.contigs.good.unique.good.filter.count_table, diffs=2)

chimera.vsearch(fasta=mapping.trim.contigs.good.unique.good.filter.unique.precluster.fasta, count=mapping.trim.contigs.good.unique.good.filter.unique.precluster.count_table, dereplicate=t)

remove.seqs(fasta=mapping.trim.contigs.good.unique.good.filter.unique.precluster.fasta, accnos=mapping.trim.contigs.good.unique.good.filter.unique.precluster.denovo.vsearch.accnos)

classify.seqs(fasta=mapping.trim.contigs.good.unique.good.filter.unique.precluster.pick.fasta, count=mapping.trim.contigs.good.unique.good.filter.unique.precluster.denovo.vsearch.pick.count_table, reference=silva.nr_v132.align, taxonomy=silva.nr_v132.tax, cutoff=80)

summary.tax(taxonomy=mapping.trim.contigs.good.unique.good.filter.unique.precluster.pick.nr_v132.wang.taxonomy, count=mapping.trim.contigs.good.unique.good.filter.unique.precluster.denovo.vsearch.pick.count_table)

dist.seqs(fasta=mapping.trim.contigs.good.unique.good.filter.unique.precluster.pick.fasta, cutoff=0.03)

cluster(column=mapping.trim.contigs.good.unique.good.filter.unique.precluster.pick.dist, count=mapping.trim.contigs.good.unique.good.filter.unique.precluster.denovo.vsearch.pick.count_table)

make.shared(list=mapping.trim.contigs.good.unique.good.filter.unique.precluster.pick.opti_mcc.list, count=mapping.trim.contigs.good.unique.good.filter.unique.precluster.denovo.vsearch.pick.count_table, label=0.03)

classify.otu(list=mapping.trim.contigs.good.unique.good.filter.unique.precluster.pick.opti_mcc.list, count=mapping.trim.contigs.good.unique.good.filter.unique.precluster.denovo.vsearch.pick.count_table, taxonomy=mapping.trim.contigs.good.unique.good.filter.unique.precluster.pick.nr_v132.wang.taxonomy, label=0.03)

rename.file(taxonomy=mapping.trim.contigs.good.unique.good.filter.unique.precluster.pick.opti_mcc.0.03.cons.taxonomy, shared=mapping.trim.contigs.good.unique.good.filter.unique.precluster.pick.opti_mcc.shared)

count.groups(shared=mapping.opti_mcc.shared)

rarefaction.single(shared=mapping.opti_mcc.shared, calc=sobs, freq=100)

summary.single(shared=mapping.opti_mcc.shared, calc=nseqs-coverage-sobs-invsimpson)

Then, mothur was performed running the command:

/mnt/big/mothur-1.42.1/mothur mothur.batch

***LotuS commands and parameters***

The sdm_miSeq.txt file was set as such:

```
#sdm options file to control sequence quality filtering, demultiplexing and preparation (can also be used without demultiplexing)
#* indicates alternative quality filtering options, saved in *.add.fna etc. files separately from initial quality filtered dataset
#sequence length refers to sequence length AFTER removal of Primers, Barcodes and trimming. this ensures that downstream analyis tools will have appropiate sequence information
#options with a star in front are lenient parameters for mid qual sequences (only used for estimating OTU abundance, not for OTU building itself).
minSeqLength	150
maxSeqLength	350
minAvgQuality	30
*minSeqLength	150
*minAvgQuality	30
#truncate total Sequence length to X (length after Barcode, Adapter and Primer removals, set to -1 to deactivate)
TruncateSequenceLength	-1

#Ambiguous bases in Sequence
maxAmbiguousNT	1
*maxAmbiguousNT	1

#sequence is discarded if a homonucleotide run in sequence is longer
maxHomonucleotide	8

#Filter whole sequence if one window of quality scores is below average
QualWindowWidth	50
QualWindowThreshhold	25

#Trim the end of a sequence if a window falls below quality threshhold. Useful for removing low qulaity trailing ends of sequence
TrimWindowWidth	20
TrimWindowThreshhold	25

#Probabilistic max number of accumulated sequencing errors. After this length, the rest of the sequence will be deleted. Complimentary to TrimWindowThreshhold. (-1) deactivates this option.
maxAccumulatedError	0.75
*maxAccumulatedError	-1
#Binomial error model of expected errors per sequence (see https://github.com/fpusan/moira), to deactivate, set BinErrorModelAlpha to -1
BinErrorModelMaxExpError	2.5
BinErrorModelAlpha	-1


#Max Barcode Errors
maxBarcodeErrs	0
maxPrimerErrs	0

#keep Barcode / Primer Sequence in the output fasta file - in a normal 16S analysis this should be deactivated (0) for Barcode and de-activated (0) for primer
keepBarcodeSeq	0
keepPrimerSeq	0


#set fastqVersion to 1 if you use Sanger, Illumina 1.8+ or NCBI SRA files. Set fastqVersion to 2, if you use Illumina 1.3+ - 1.7+ or Solexa fastq files.  "auto" will look for typical characteristics of either of these and choose the quality offset score automatically.
fastqVersion	auto

#if one or more files have a technical adapter still included (e.g. TCAG 454) this can be removed by setting this option
TechnicalAdapter

#delete X NTs (e.g. if the first 5 bases are known to have strange biases)
TrimStartNTs	0

#correct PE header format (1/2) this is to accomodate the illumina miSeq paired end annotations 2="@XXX 1:0:4" insteand of 1="@XXX/1". Note that the format will be automatically detected
PEheaderPairFmt	1

#sets if sequences without match to reverse primer (ReversePrimer) will be accepted (T=reject ; F=accept all); default=F
RejectSeqWithoutRevPrim	F
#*RejectSeqWithoutRevPrim	F
#sets if sequences without a forward (LinkerPrimerSequence) primer will be accepted (T=reject ; F=accept all); default=F
RejectSeqWithoutFwdPrim	F
#*RejectSeqWithoutFwdPrim	F

#this option should be "T" if your amplicons are possibly shorter than a single read in a paired end sequencing run (e.g. if the 16S amplicon length is 200bp in a 250x2 miSeq run, set this to "T"). This option increases runtime by 10%, if in doubt just set to "T". *Requires* LinkerPrimerSequence and ReversePrimer to be defined in mapping file.
AmpliconShortPE	F

#options for difficulties during sequencing library construction
#checks if pair1 and pair2 were switched (ignore if single read data)
CheckForMixedPairs	F
#checks if whole amplicon was reverse-transcribed sequenced (not switched, just reverse translated)
CheckForReversedSeqs	F
```

Then, LotuS was performed running the command:

Perl /home1/haris/lotus_pipeline/lotus.pl -i /home1/haris/MyData/ -o /home1/haris/MyData_LotuS/output/ -m /home1/haris/MyData_LotuS/mymap.txt -c /home1/haris/lotus_pipeline/lOTUs.cfg -s /home1/haris/MyData_LotuS/sdm_miSeq.txt -keepTmpFiles 1 -highmem 1 -p miSeq -itsextraction 0 -simBasedTaxo 2 -refDB SLV

***PEMA command and parameters***

The parameters file for the case of the 16S rRNA marker gene and the [30] dataset was set as such:

```
outputFolderName	amvrakikos_silva132_final
EnaData	Yes
maxInfo	Yes
targetLength	200
strictness	0.3
adapters	TruSeq2-PE.fa
seedMismatches	0
palindromeClipThreshold	20
simpleClipThreshold	30
leading 	20
trailing		20
minlen	100
threadsTrimmomatic	20
pandaseqAlgorithm	simple_bayesian
vsearchThreads	20
vsearchId	0.98
gene	gene_16S
taxonomyAssignmentMethod	alignment
silvaVersion	silva_132
taxonomyFolderName	16S_taxon_assign
phyloseq	No
raxmlThreads	20
parsTrees	10
bootstrapTrees	100
emptyRawDataFile	Yes
emptyCheckpoints	Yes
```

Then, PEMA was performed running the command:

singularity run -B /home1/haris/16S_analysis/:/mnt /home1/haris/metabar_pipeline/pema.simg

Similarly, for the case of the COI marker gene and the [31] dataset, the parameters were the following:

```
outputFolderName	lakes_235bp_only_d_10
EnaData	Yes
targetLength	235
strictness	0.4
adapters	TruSeq2-PE.fa
seedMismatches	1
palindromeClipThreshold	20
simpleClipThreshold	30
leading	20
trailing	20
minlen	100
spadesAlgorithm	simple_bayesian
vsearchThreads	20
clusteringAlgo	algo_SWARM
d	10
abskew	2
emptyRawDataFile	No
emptyCheckpoints	Yes
```

For the rest of the runs that are presented in this study, the only changes in the parameter set above, was the value of the cluster radius (*d*) of the Swarm algorithm. Obviously, the “outputFolderName” was also changed correspondingly in each run.
