## Supplementary material for "PEMA: from the raw .fastq files of 16S rRNA and COI marker genes to the (M)OTU-table, a thorough metabarcoding analysis": Table S1

Table S1: Number of sequences after each pre-processing step for the case of COI dataset from [31].

| **sample libraries** | **run_accession** | **initial number of reads** | **after trimming** | **after specific-position error correction (Bayes-Hammer/ Spades)** | **after merging (PANDAseq)** | **after dereplication (obiuniq)** |
| --- | --- | --- | --- | --- | --- | --- |
| 1_WCOI_S | ERR1308201 | 284052 | 246323 | 246318 | 16633 | 15171 |
| 2_WCOI_S | ERR1308202 | 231471 | 213037 | 213033 | 9177 | 8781 |
| 3_WCOI_S | ERR1308203 | 99362 | 68316 | 68302 | 3492 | 3458 |
| 4_WCOI_S | ERR1308204 | 335506 | 312552 | 312546 | 16610 | 15324 |
| 5_WCOI_S | ERR1308205 | 313499 | 279436 | 279431 | 9955 | 9477 |
| 6_WCOI_S | ERR1308206 | 328491 | 313222 | 313215 | 15446 | 13036 |
| 7_WCOI_S | ERR1308207 | 193450 | 171005 | 170999 | 10593 | 9945 |
| 8_WCOI_S | ERR1308208 | 308713 | 286960 | 286956 | 14475 | 13049 |
| 9_WCOI_S | ERR1308209 | 371277 | 357721 | 357713 | 16440 | 14948 |
| 9_ECOI_S | ERR1308210 | 711 | 574 | 571 | 29 | 29 |
| 2_ECOI_S | ERR1308211 | 710 | 585 | 582 | 37 | 37 |
| 10_WCOI_S | ERR1308233 | 332045 | 317014 | 317008 | 14074 | 12486 |
| 11_WCOI_S | ERR1308234 | 267676 | 254660 | 254654 | 11091 | 10418 |
| 12_WCOI_S | ERR1308235 | 330533 | 313003 | 312997 | 17041 | 16477 |
| 13_WCOI_S | ERR1308236 | 297504 | 282663 | 282655 | 16034 | 14999 |
| 14_WCOI_S | ERR1308237 | 329782 | 320270 | 320265 | 15419 | 13429 |
| 15_WCOI_S | ERR1308238 | 567252 | 507855 | 507834 | 42907 | 25635 |
| 16_WCOI_S | ERR1308239 | 370273 | 353640 | 353632 | 18843 | 17606 |
| 3_ECOI_S | ERR1308240 | 1323 | 529 | 526 | 20 | 20 |
| 4_ECOI_S | ERR1308241 | 595 | 428 | 425 | 15 | 15 |
| 5_ECOI_S | ERR1308242 | 248774 | 201093 | 201089 | 9584 | 6487 |
| 6_ECOI_S | ERR1308243 | 873 | 418 | 416 | 26 | 26 |
| 7_ECOI_S | ERR1308244 | 787 | 299 | 295 | 8 | 8 |
| 8_ECOI_S | ERR1308245 | 631 | 470 | 468 | 18 | 18 |
| 1_ECOI_S | ERR1308246 | 191597 | 163621 | 163615 | 9629 | 8832 |
| 10_ECOI_S | ERR1308247 | 196460 | 167200 | 167191 | 10999 | 9755 |
| 11_ECOI_S | ERR1308248 | 166685 | 150463 | 150457 | 8482 | 7685 |
| 12_ECOI_S | ERR1308249 | 366491 | 285717 | 285715 | 11649 | 10627 |
| 13_ECOI_S | ERR1308250 | 195820 | 167381 | 167375 | 8728 | 7885 |
| 14_ECOI_S | ERR1308251 | 436253 | 357580 | 357574 | 22389 | 19617 |
| 15_ECOI_S | ERR1308252 | 388763 | 312722 | 312718 | 24696 | 19640 |
| 16_ECOI_S | ERR1308253 | 624992 | 504707 | 504698 | 28796 | 25970 |
| **SUM** |  | **7782351** | **6911464** | **6911273** | **383335** | **330890** |
