## Supplementary material for "PEMA: from the raw .fastq files of 16S rRNA and COI marker genes to the (M)OTU-table, a thorough metabarcoding analysis": Table S2

Table S2: Number of sequences after each pre-processing step for the case of 16S rRNA gene.

| **sample libraries** | **run_accession** | **initial number**  **of reads** | **after**  **trimming** | **after specific-position**  **error correction**  **(Bayes-Hammer/ Spades)** | **after merging**  **(PANDAseq)** | **after dereplication**  **(obiuniq)** |
| --- | --- | --- | --- | --- | --- | --- |
| L_LOout_A | ERR1906853 | 96457 | 95748 | 95569 | 46576 | 37503 |
| L_LOout_B | ERR1906854 | 144529 | 142133 | 141847 | 65556 | 56636 |
| L_LOout_C | ERR1906855 | 139149 | 138811 | 138490 | 64650 | 51889 |
| L_LOin_A | ERR1906856 | 130200 | 128734 | 128462 | 59521 | 51640 |
| L_LOin_B | ERR1906857 | 108906 | 108115 | 107927 | 51067 | 41901 |
| L_LOin_C | ERR1906858 | 110955 | 109660 | 109428 | 51843 | 45147 |
| S_Kal_A | ERR1906859 | 95387 | 94836 | 94615 | 41880 | 37249 |
| S_Kal_B | ERR1906860 | 100113 | 99321 | 99087 | 44317 | 41666 |
| S_Kal_C | ERR1906861 | 100822 | 100293 | 100012 | 44011 | 40730 |
| R_ARDelta_A | ERR1906862 | 99760 | 99076 | 98734 | 34367 | 33110 |
| R_ARDelta_B | ERR1906863 | 120241 | 119374 | 118970 | 42085 | 39841 |
| R_ARDelta_C | ERR1906864 | 94144 | 93618 | 93131 | 32188 | 30936 |
| R_AR_A | ERR1906865 | 97520 | 96530 | 96379 | 46297 | 41914 |
| R_AR_B | ERR1906866 | 157246 | 156223 | 155950 | 76214 | 70533 |
| R_AR_C | ERR1906867 | 167487 | 167016 | 166742 | 80555 | 73018 |
| R_ARO_A | ERR1906868 | 124486 | 123858 | 123648 | 54574 | 52558 |
| R_ARO_B | ERR1906869 | 148083 | 147656 | 147402 | 68979 | 62077 |
| R_ARO_C | ERR1906870 | 142040 | 141718 | 141455 | 65768 | 58984 |
| **SUM** |  | **4355050** | **4325440** | **4315704** | **1036216** | **867332** |

All sample libraries that start with “L” correspond to samples collected from lagoons, with “S” from the sea while “R” stands for the riverine samples. AR: Arachthos. ARO: Arachthos Neochori. ARDelta: Arachthos Delta. LOin: Logarou station inside the lagoon. LOout: Logarou station in the channel connecting the lagoon to the gulf. Kal: Kalamitsi. A, B, C replicate samples.
