## Supplementary figures and images for "PEMA: from the raw .fastq files of 16S rRNA and COI marker genes to the (M)OTU-table, a thorough metabarcoding analysis"

### Figure S1

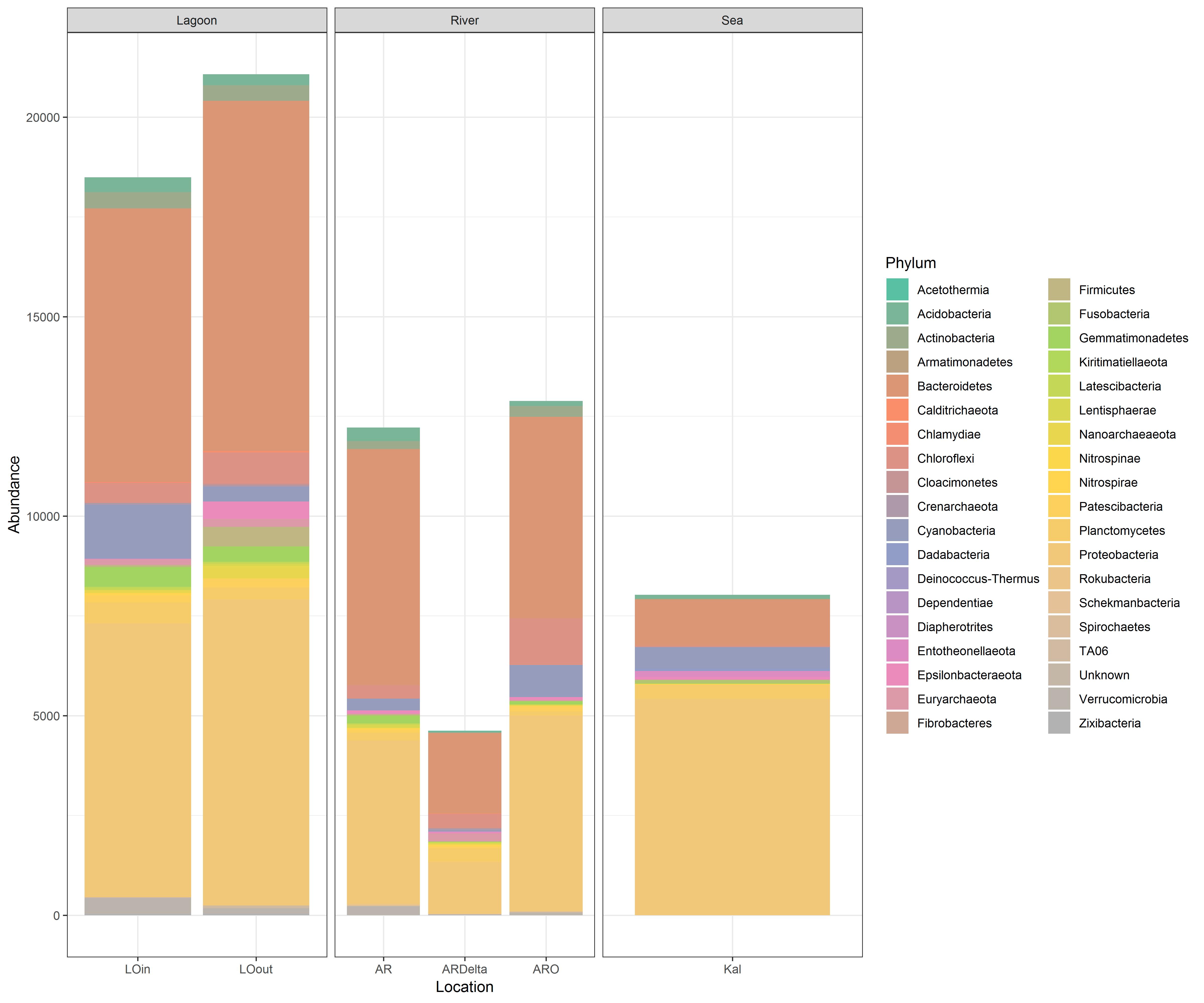

### Figure S3

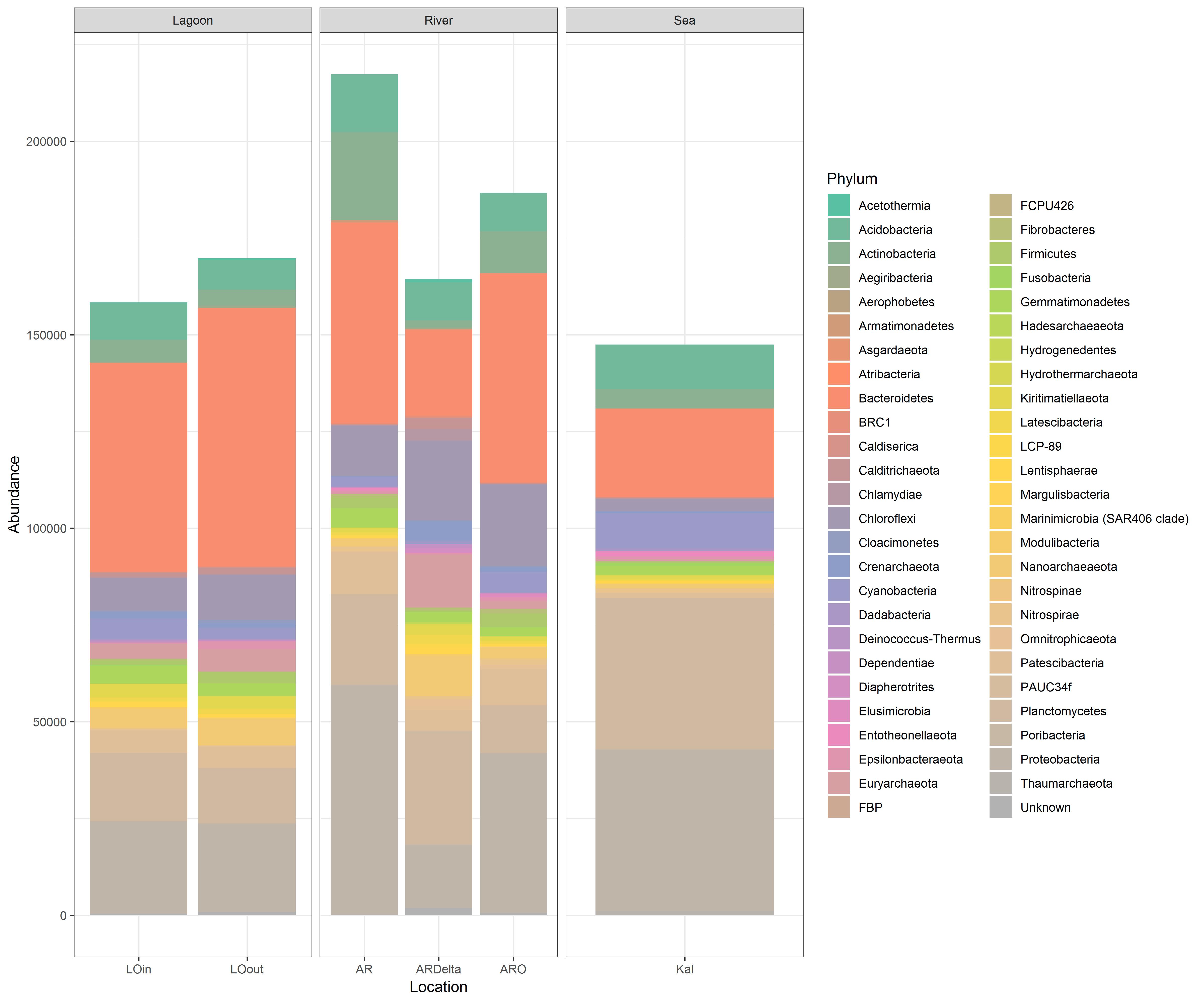

### Figure S4

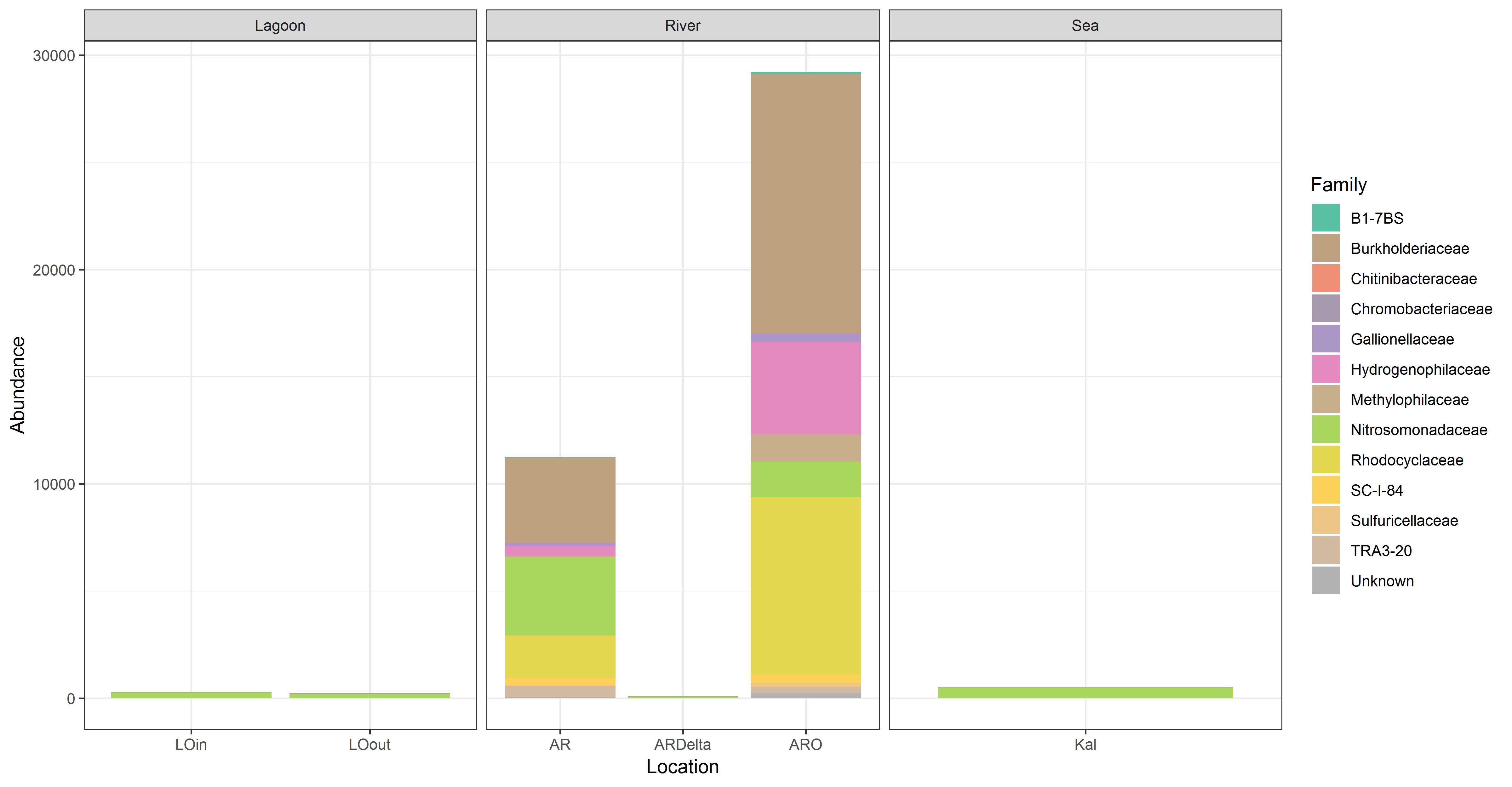

### Figure S5

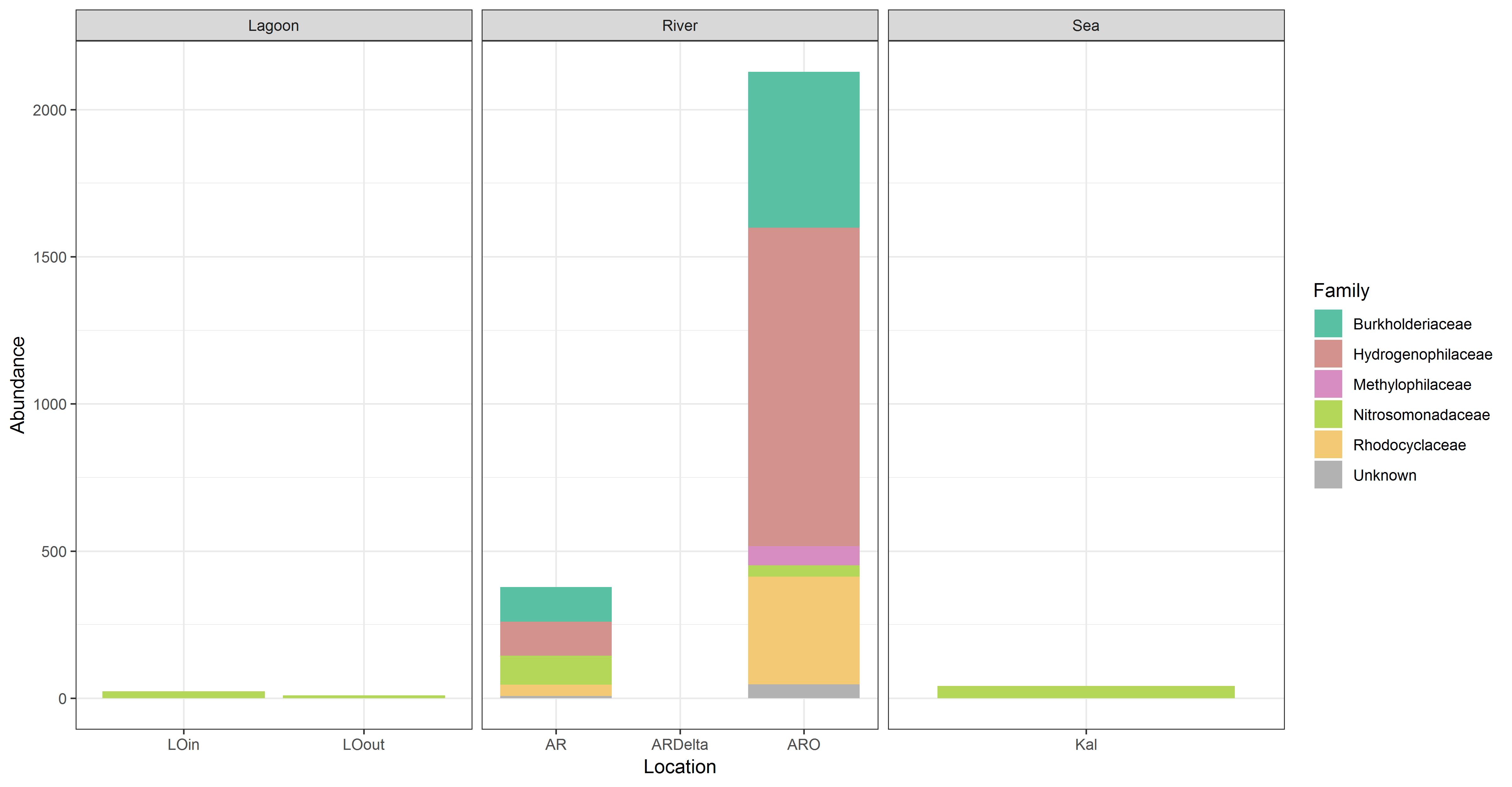

### Figure S6

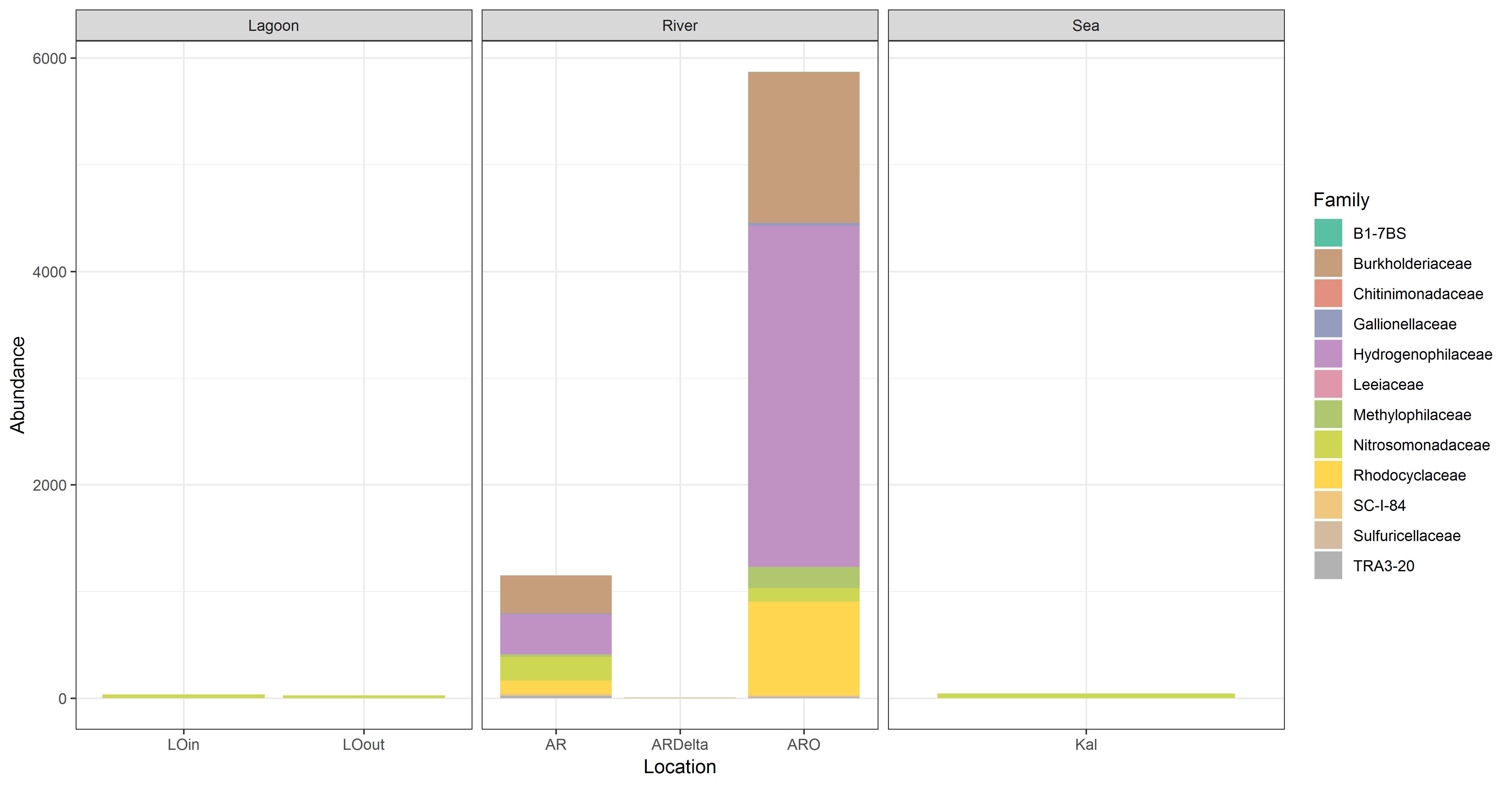
